# Altered spike timing-dependent plasticity rules in physiological calcium

**DOI:** 10.1101/2020.03.16.993675

**Authors:** Yanis Inglebert, Johnatan Aljadeff, Nicolas Brunel, Dominique Debanne

## Abstract

Like many forms of long-term synaptic plasticity, spike-timing-dependent plasticity (STDP) depends on intracellular Ca^2+^ signaling for its induction. Yet, all *in vitro* studies devoted to STDP used abnormally high external Ca^2+^ concentration. We measured STDP at the CA3-CA1 hippocampal synapses under different extracellular Ca^2+^ concentrations and found that the sign, shape and magnitude of plasticity strongly depend on Ca^2+^. A pre-post protocol that results in robust LTP in high Ca^2+^, yielded only LTD or no plasticity in the physiological Ca^2+^ range. LTP could be restored by either increasing the number of post-synaptic spikes or increasing the pairing frequency. A calcium-based plasticity model in which depression and potentiation depend on post-synaptic Ca^2+^ transients was found to fit quantitatively all the data, provided NMDA receptor-mediated non-linearities were implemented. In conclusion, STDP rule is profoundly altered in physiological Ca^2+^ but specific activity regimes restore a classical STDP profile.

## Introduction

Spike-timing-dependent plasticity (STDP) is a form of synaptic modification that is controlled by the relative timing between pre- and post-synaptic activity and depends on intracellular Ca^2+^ signaling (review in ^1, 2^). Timing-dependent long-term synaptic potentiation (t-LTP) is induced when synaptic activity is followed by one or more backpropagating action potentials in the post-synaptic cell ^3–8^. It involves post-synaptic Ca^2+^ influx through NMDA receptors that in turn activates protein kinases ^3, 6, 8, 9^. Timing-dependent long-term synaptic depression (t-LTD) is expressed when synaptic activity is repeatedly preceded by one or more backpropagating action potentials ^4–7, 10^. It depends on NMDA receptor activation, post-synaptic metabotropic glutamate receptors (mGluR), voltage-dependent calcium channels, protein phosphatases, cannabinoid receptor CB1 and astrocytic signaling ^6, 10–16^. Calcium therefore represents potentially a key factor in the induction of STDP. The intracellular Ca^2+^ dependence of STDP suggests that extracellular Ca^2+^ might play a critical role in shaping STDP. Yet, most if not all *in vitro* STDP studies ^6–10, 17–19^ used non-physiological external Ca^2+^ concentrations ranging between 2 and 3 mM, whereas the physiological Ca^2+^ concentration ranges between 1.3 and 1.8 mM in the young rat hippocampus ^20, 21^. Calcium-based models of synaptic plasticity ^22, 23^ where Ca^2+^ transients result from backpropagating action potentials and EPSPs predict that the sign, shape and magnitude of STDP strongly depend on the amplitudes of calcium transients triggered by pre and post synaptic spikes, and therefore on external Ca^2+^ concentration ^23^ (see Figure 1). These modeling studies suggest that high Ca^2+^ concentrations used in experimental studies could lead to an overestimate of the *in vivo* levels of plasticity, or even to a lack of plasticity. We therefore set out to determine STDP rules in physiological Ca^2+^ at the CA3-CA1 synapse of the hippocampus *in vitro*.

**Figure 1.**
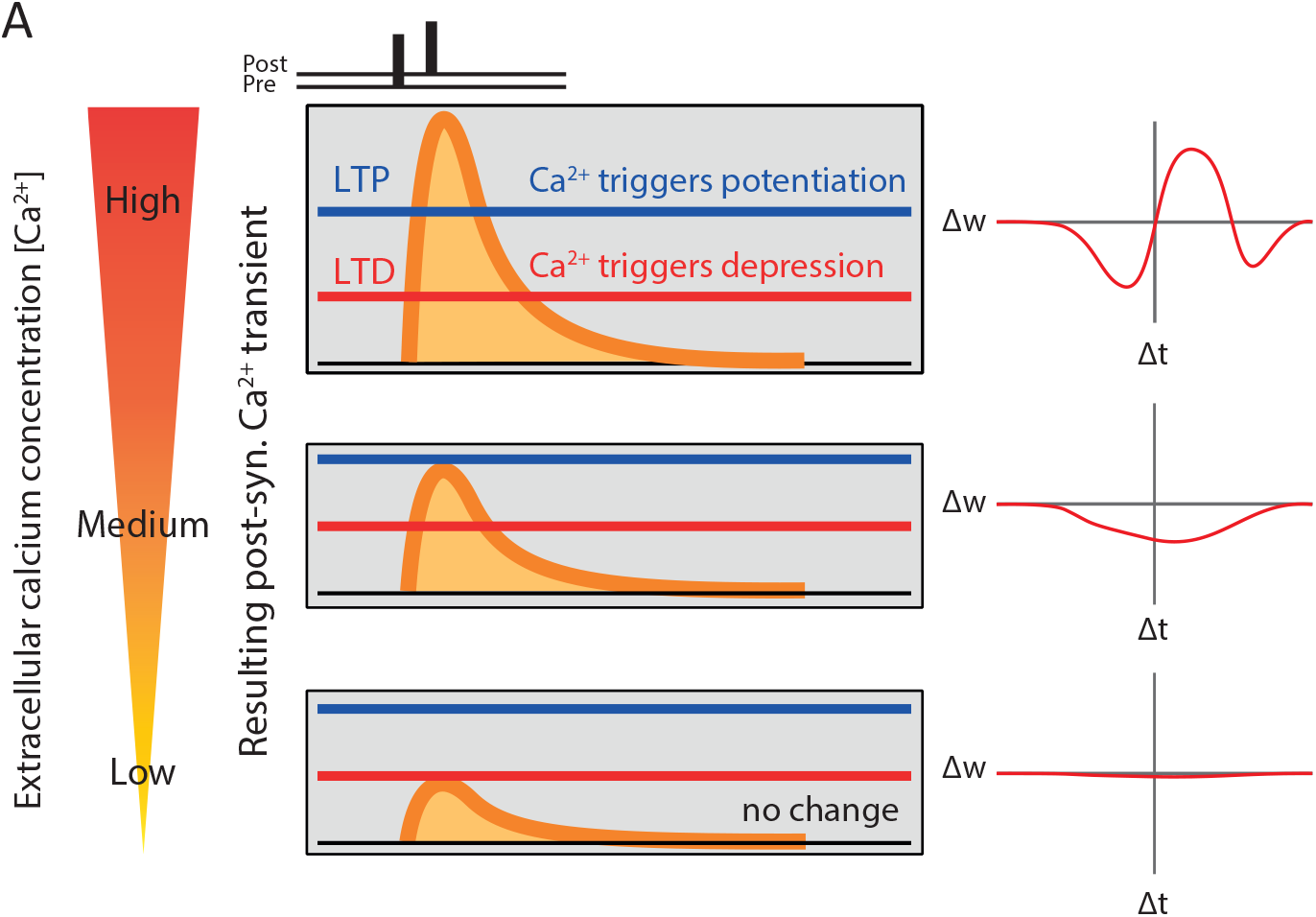
Calcium-based model of Spike-Timing-Dependent Plasticity. Cartoon qualitatively showing that a calcium-based model of synaptic plasticity implies that the shape of STDP (and sign of plasticity) will depend on extracellular calcium concentration. Synaptic changes depend on two plasticity thresholds, one for LTP (blue) and one for LTD (red). In response to a pre-post protocol, the post-synaptic calcium transient determines the orientation of plasticity. The reduction of extracellular calcium reduces the post-synaptic calcium transient, and leads to LTP for average values and an absence of plasticity for low values.

We show here that the classical STDP rule (LTD for post-before-pre pairings, LTP for pre-before-post pairings) is obtained solely with a high external Ca^2+^ concentration (≥ 2.5 mM), whereas no plasticity could be induced for concentrations lower than 1.5 mM external Ca^2+^, and only t-LTD could be induced by positive or negative time delays in 1.8 mM external Ca^2+^. t-LTP could be restored only when bursts of 3 or 4 post-synaptic spikes were used instead of single spikes, or when the pairing frequency was increased from 0.33 to 5 or 10 Hz. We used two variants of a Ca^2+^-based plasticity model ^23^ in which both t-LTD and t-LTP depend on transient changes in post-synaptic Ca^2+^ (Figure 1) to fit the data. We found that the nonlinearity of transient Ca^2+^ changes conferred by NMDA receptor activation is critical to quantitatively account for the entire experimental dataset. Our results indicate that the STDP rule is profoundly altered in physiological Ca^2+^, but that a classical STDP profile can be restored under specific activity regimes.

## Results

### STDP rule is altered in extracellular physiological calcium

Induction of STDP at the Schaffer collateral-CA1 cell synapses was examined under 3 different external Ca^2+^ concentrations (3, 1.8 and 1.3 mM) and in the presence of normal synaptic inhibition (i.e. without any GABA_A_ receptor antagonist). In high Ca^2+^ (3 mM), t-LTP (124 ± 7% of the control EPSP slope, n = 14) was induced by repeatedly pairing (100 times, 0.3 Hz ^24^; Figure 2A) the evoked excitatory post-synaptic potential (EPSP) and the post-synaptic action potential (AP) with a delay ranging between +5 and +25 ms (Figure 2B). t-LTD (68 ± 11%, n = 10) was induced by repeatedly pairing (150 times, 0.3 Hz ^24^; Figure 2A) the EPSP and the post-synaptic AP with a delay ranging between −5 and −25 ms (Figure 2B). The plot of synaptic changes as a function of spike-timing provided a classical STDP curve in 3 mM external Ca^2+^, with a t-LTP window in the [0,40ms] range surrounded by two t-LTD windows, one for negative delays and the other for positive delays at ∼40-60 ms (Figure 2A). In contrast, only t-LTD was induced by positive or negative pairings in 1.8 mM external Ca^2+^ (respectively, 73 ± 6%, n = 13 & 71 ± 8%, n=11; Figure 2C) and no significant changes were observed in 1.3 mM external Ca^2+^ at any pairing (100 ± 12%, n = 13 after positive pairing and 106 ± 14%, n = 13 after negative pairing; Figure 2D). Thus, the STDP rules are highly altered in the physiological Ca^2+^ range (Figure 2C & 2D), as predicted by calcium based models ^22, 23^ (Fig.1).

**Figure 2.**
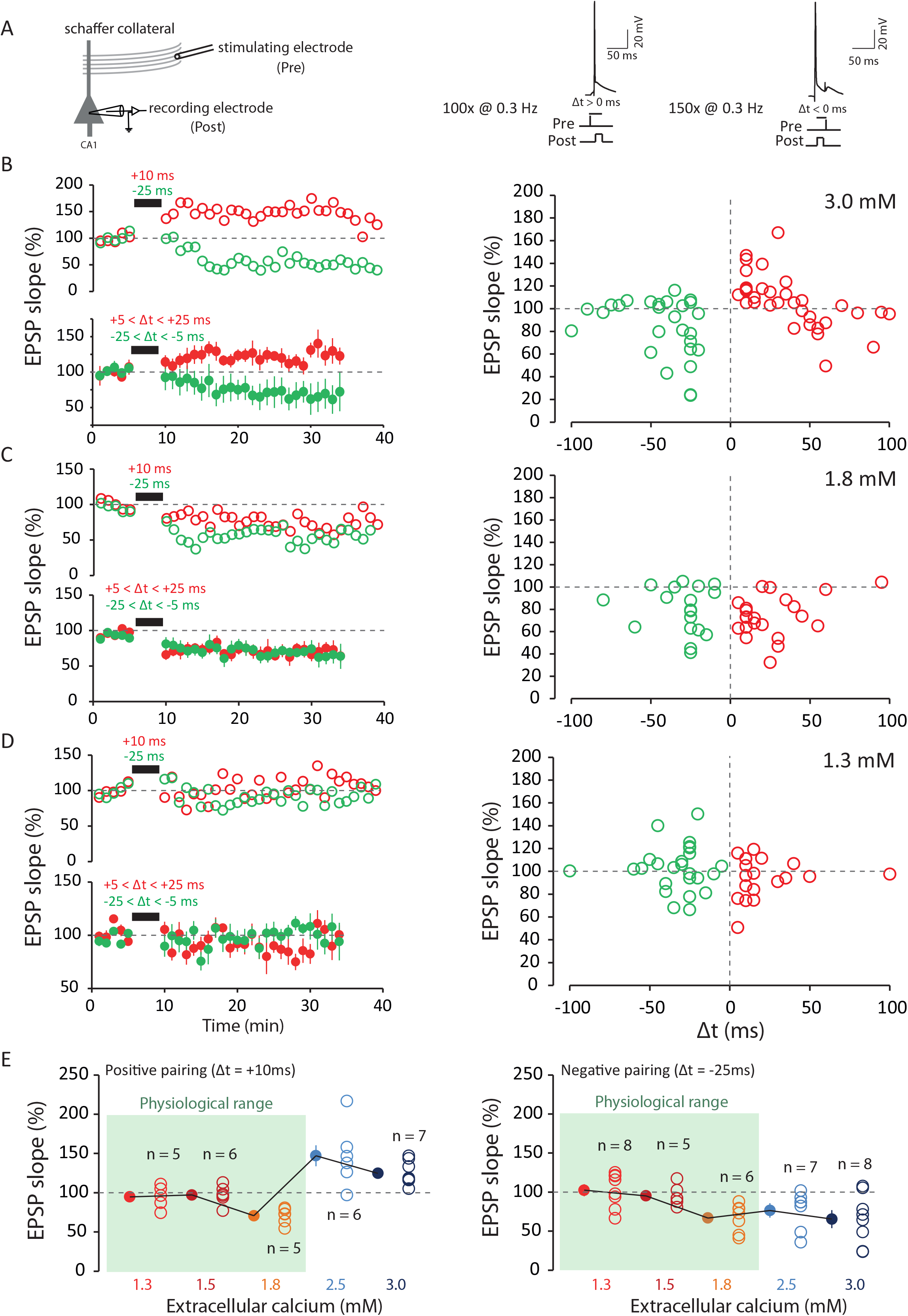
STDP under various external calcium concentration. **(A)** t-LTP and t-LTD are induced at low frequency (0.3 Hz). The pre-post protocol is repeated 100 times while the post-pre-post protocol is repeated 150 times. The action potential is induced by the injection of a depolarizing current into a CA1 neuron recorded in a whole cell configuration. The PPSE is induced by a stimulation electrode placed in the Schaffer collaterals. The delay (Δt) is varied according to the experiments. **(B)** In 3 mM extracellular calcium, a pre-post protocol (positive delays; +5 < Δt < +25 ms; red) leads to t-LTP and a post-pre (negative delays; −25 < Δt < −5 ms; green) protocol leads to t-LTD. Note the presence of a second t-LTD window at around +40 / +60 ms. **(C)** In 1.8 mM extracellular calcium, a pre-post protocol (+5 < Δt < +25 ms; red) leads to t-LTD while a post-pre protocol (−25 < Δt < −5 ms; green leads to t-LTD. The t-LTP window is absent under these conditions. **(D)** In 1.3 mM extracellular calcium, on average, no plasticity is induced regardless of the delay. The t-LTP and t-LTD window are missing under these conditions. **(E)** Left, synaptic changes for a pre-post protocol at +10 ms. No plasticity is induced for calcium concentrations of 1.3 and 1.5 mM. For 1.8, 2.5 and 3 mM calcium, t-LTP is induced. Right, synaptic changes for a post-pre protocol at −25 ms. No plasticity is induced in 1.3 and 1.5 mM calcium. For 1.8, 2.5 and 3 mM calcium, a t-LTD is induced. Note the difference between the results in the range of physiological calcium concentration (green squares) and the results in non-physiological calcium concentration.

In order to precisely determine the transition between t-LTD and no change on the negative correlation side, and the transition from t-LTP to t-LTD to no change on the positive correlation side, we tested the effects of positive and negative pairing in 1.5 mM and 2.5 mM external Ca^2+^. The data show that negative pairing induced no plasticity in 1.5 mM external Ca^2+^ (95 ± 5%, n = 5, Figure 2E), whereas t-LTD was consistently observed in the 1.8-3 mM range (at 2.5 mM, 90 ± 3%, n = 5, Figure 2E). Positive pairing led to no change in 1.5 mM external Ca^2+^ (97 ± 5%, n = 6), t-LTD at 1.8 mM external Ca^2+^, but t-LTP in 2.5 mM external Ca^2+^ (147 ± 13%, n = 6), with a switch from t-LTD to t-LTP occurring between 1.8 and 2.5 mM (Figure 2E). These data indicate that the STDP rule is profoundly altered in physiological calcium, with a lack of t-LTP in the whole range, and the appearance of t-LTD in a broad range of timings in the upper limit of the physiological Ca^2+^ range.

### Calcium imaging

In order to better understand why no plasticity occurred in 1.3 mM, calcium transients in response to positive (+20 ms) and negative pairings (−20 ms) between an evoked EPSP and a post-synaptic spike, were measured using Fluo-4 in dendritic spines (Figure 3A). While the magnitude of the calcium transients was found to be much greater for positive pairing than for negative pairing in 3 mM calcium (ratio PP/NP = 250 ± 27%, n = 7; Figure 3B & C), the difference in calcium transient amplitude were found to be considerably smaller in 1.3 mM (ratio PP/NP = 132 ± 17%, n = 6, MW p<0.01; Figure 3B & 3C). Thus, the lack of plasticity in 1.3 mM may result from reduced calcium transient amplitudes.

**Figure 3.**
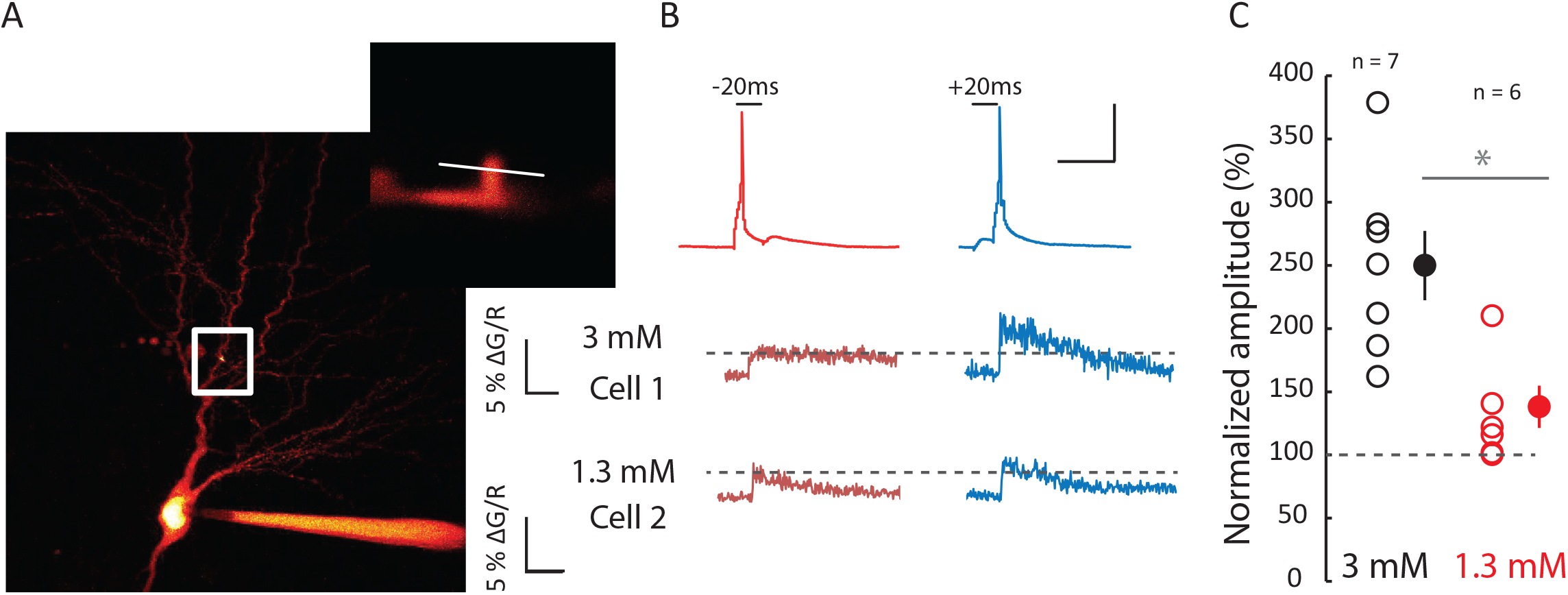
Calcium imaging. **(A)** Comparison of calcium spine entry during negative and positive pairing in 3 mM and 1.3 mM extracellular calcium. Calcium was measured in the spine of a CA1 pyramidal cell loaded with 50 µM Fluo-4. (**B)** Each neuron was alternatively test for negative (Δt = −20 ms) and positive pairing (Δt = +20 ms). Cell 1 recorded in 3 mM calcium displays a much larger influx of calcium in response to positive pairing than negative pairing. Cell 2 recorded in 1.3 mM calcium displays an almost equal calcium rise. (**C**) Pooled data of the normalized ratio of calcium evoked by positive pairing over negative pairing (p<0.01 Mann-Whitney U-test).

### Recovery of t-LTP in physiological calcium

The STDP rule in physiological Ca^2+^ is characterized by a complete lack of t-LTP and the predominance of t-LTD at 1.8 mM external Ca^2+^, while no significant plasticity is observed at 1.3 mM. Therefore, we first tried to restore t-LTP in physiological Ca^2+^ by either using bursts of post-synaptic spikes, instead of individual spikes, during the pairing ^18, 25^ or increasing the frequency of single-spike pairing ^18^. In 1.8 mM external Ca^2+^, increasing the number of post-synaptic spikes in a burst from 1 to 4 increased almost linearly the magnitude of potentiation as a function of the number of spikes per burst. While no net change was observed with 2 spikes (99 ± 15%, n = 11), significant potentiation was observed with 3 (116 ± 6%, n = 8) and 4 spikes (135 ± 13, n = 7; Figure 4A). In contrast, no potentiation was induced with 3 postsynaptic spikes in 1.3 mM external Ca^2+^ (88 ± 7%, n = 7; Figure 4B).

**Figure 4.**
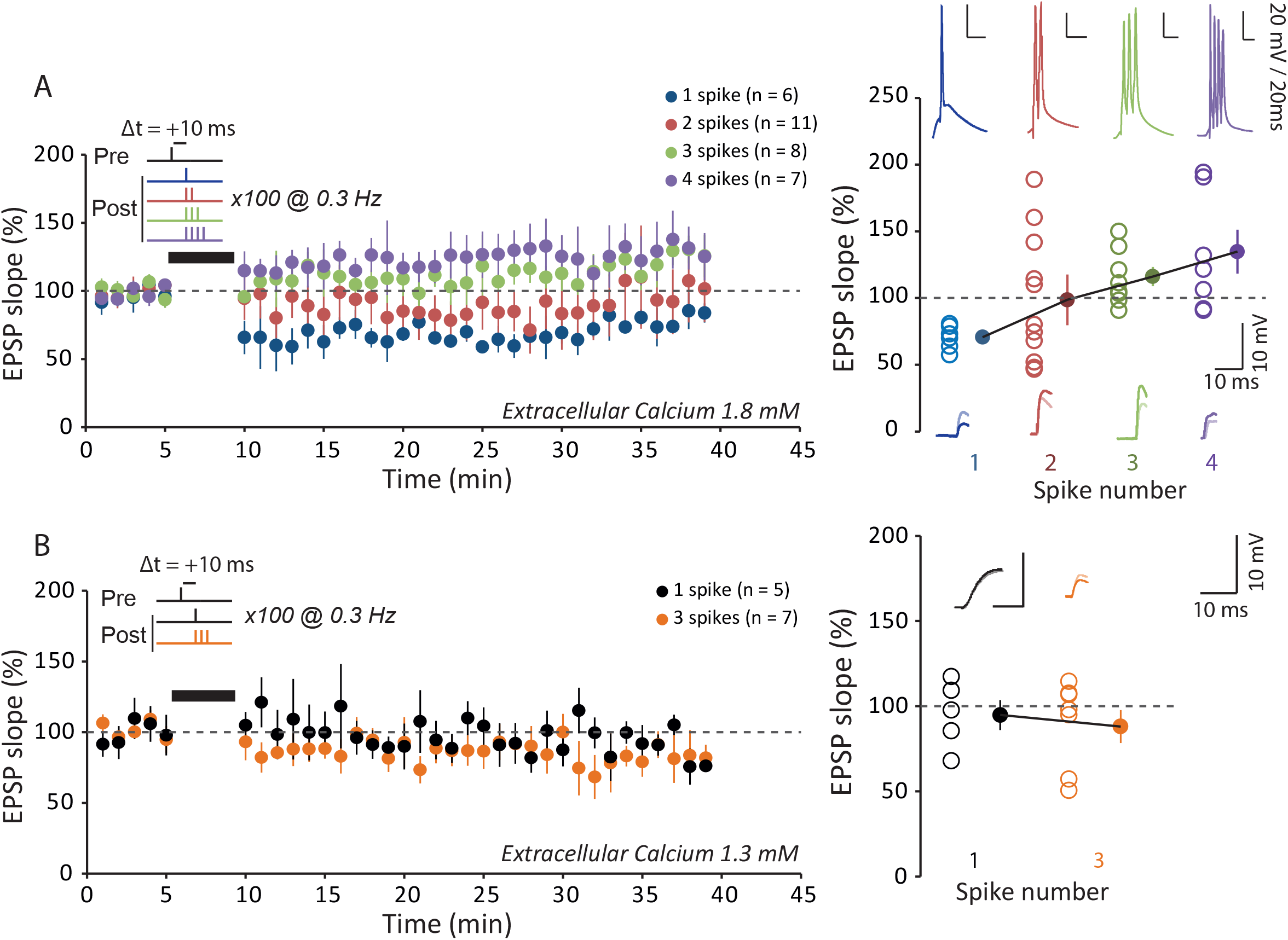
Recovery of t-LTP with increasing spike number. **(A)** Role of post-synaptic spike number for inducing synaptic changes by a pre-post protocol at +10 in 1.8 mM extracellular calcium. With only one action potential (blue), t-LTD is induced. With two action potentials (red), on average, no plasticity is induced. Note the large variability in the plasticity. With three (green) or four (purple) action potentials t-LTP is induced. The increase in the number of post-synaptic action potential allows t-LTP to be restored. **(B)** Role of post-synaptic spike number for inducing synaptic changes by a pre-post protocol at +10 in 1.3 mM extracellular calcium. With only one action potential (black), no plasticity is induced. With three action potentials (orange), t-LTP does not recover.

To understand why increasing spike number in 1.3 mM did not recover t-LTP, we performed calcium imaging in dendritic spine upon positive pairings with 1, 2, 3 and 4 spikes. While in 3 mM calcium a clear increase in calcium signal was observed when spike number was increased (for 2 spikes: 146 ± 17%, for 3 spikes: 230 ± 38% and for 4 spikes: 261 ± 49% of the calcium signal obtained for 1 spike; Supplementary Figure 1), no calcium rise was observed in 1.3 mM (for 2 spikes: 89 ± 7% and for 3 spikes: 89 ± 7% and for 4 spikes: 99 ± 4% of the calcium signal obtained for 1 spike; Supplementary Figure 1), thus confirming the fact that, in physiological conditions, calcium dynamics is considerably reduced.

We next increased the frequency of single-spike pairing from 0.5 Hz to 3-10 Hz in physiological external Ca^2+^. While no net change was observed when the frequency of stimulation was increased to 3 Hz (95 ± 4%, n = 8), significant potentiation was observed with 5 (138 ± 17%, n = 7) or 10 Hz (131 ± 9 %, n = 8) in 1.8 mM external Ca^2+^ (Figure 5A). Increasing the frequency of stimulation to 10 Hz also allowed to recover t-LTP in 1.3 mM external Ca^2+^ (129 ± 9%, n = 9, Figure 5B). We conclude that while STDP rules are strongly altered, t-LTP can be restored either by increasing the post-synaptic spike number or single-spike pairing frequency in physiological Ca^2+^.

**Figure 5.**
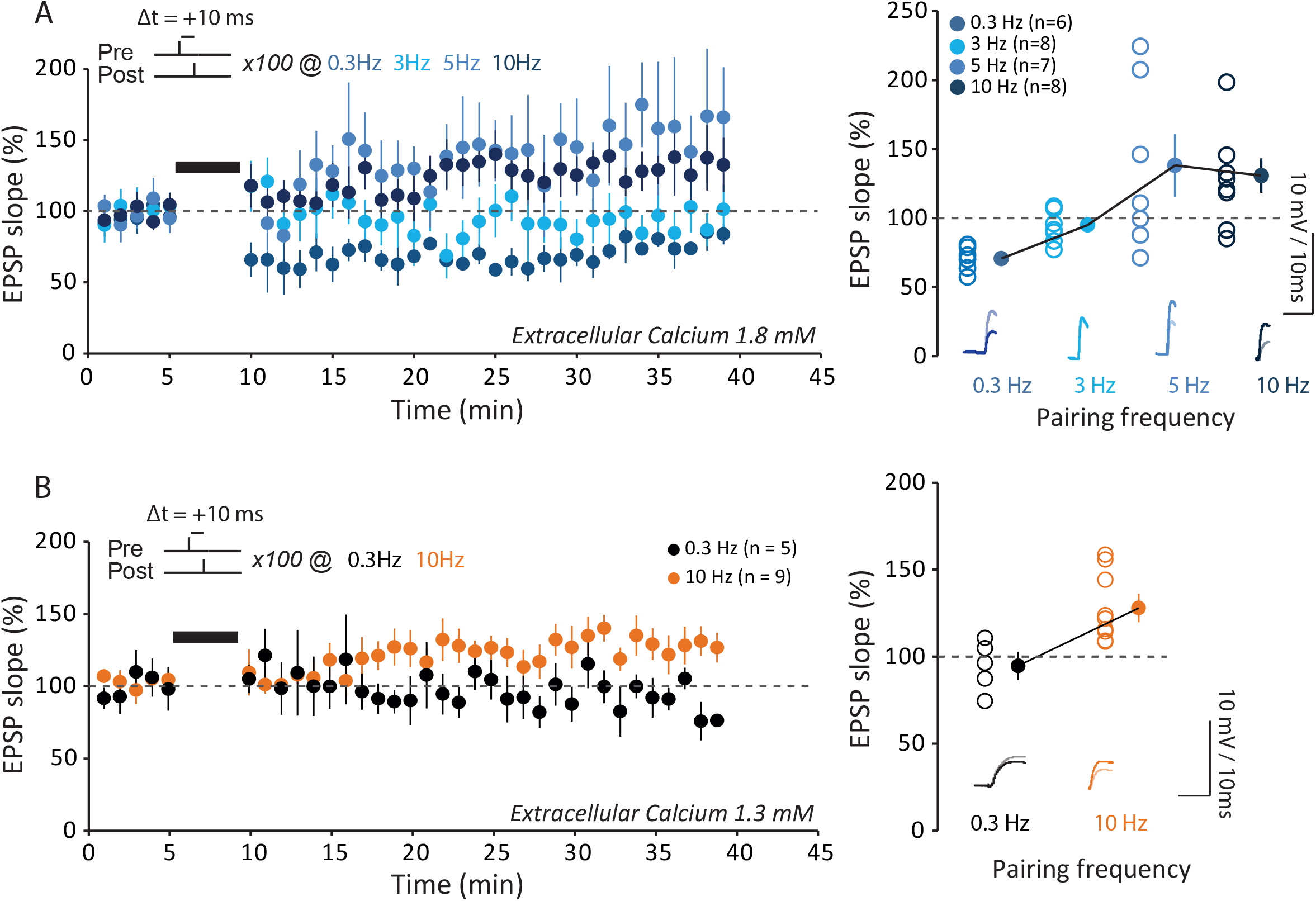
Recovery of t-LTP with increasing pairing frequency. **(A)** Effects of the stimulation frequency on synaptic changes induced by a pre-post protocol at +10 ms with a single post-synaptic action potential in 1.8 mM extracellular calcium. For 0.3 Hz, t-LTD is induced. For 3 Hz, no synaptic changes are induced. For 5 or 10 Hz, a LTP is induced. Increasing the pairing frequency restores t-LTP. **(B)** Effects of the stimulation frequency on synaptic changes induced by a pre-post protocol at +10 ms with a single post-synaptic action potential in 1.3 mM extracellular calcium. For 0.3 Hz, no synaptic change is induced. For 10 Hz, a LTP is induced. Increasing the pairing frequency restores t-LTP.

### Recovery of t-LTD at negative delays in 1.3 mM calcium

While t-LTD induced by negative delays was observed at the upper limit of the physiological calcium range (i.e. 1.8 mM), it was absent in 1.3 mM external Ca^2+^ (106 ± 14%, n = 10; Figure 6A). We therefore examined the conditions for restoring t-LTD in 1.3 mM Ca^2+^ at negative delays. When the number of post-synaptic spikes was increased up to 3 during the pairing (delay of −25 ms), significant t-LTD was found to be induced (61 ± 7%, n = 7, MW p<0.01; Figure 6A). Similarly, when the pairing frequency was increased from 0.3 to 10 Hz, significant t-LTD was restored (75 ± 6%, n = 9, MW p<0.01; Figure 6B). We conclude that normal STDP profile can be restored even in 1.3 mM Ca^2+^ by either increasing the number of post-synaptic spikes during the pairing or the pairing frequency to 10 Hz.

**Figure 6.**
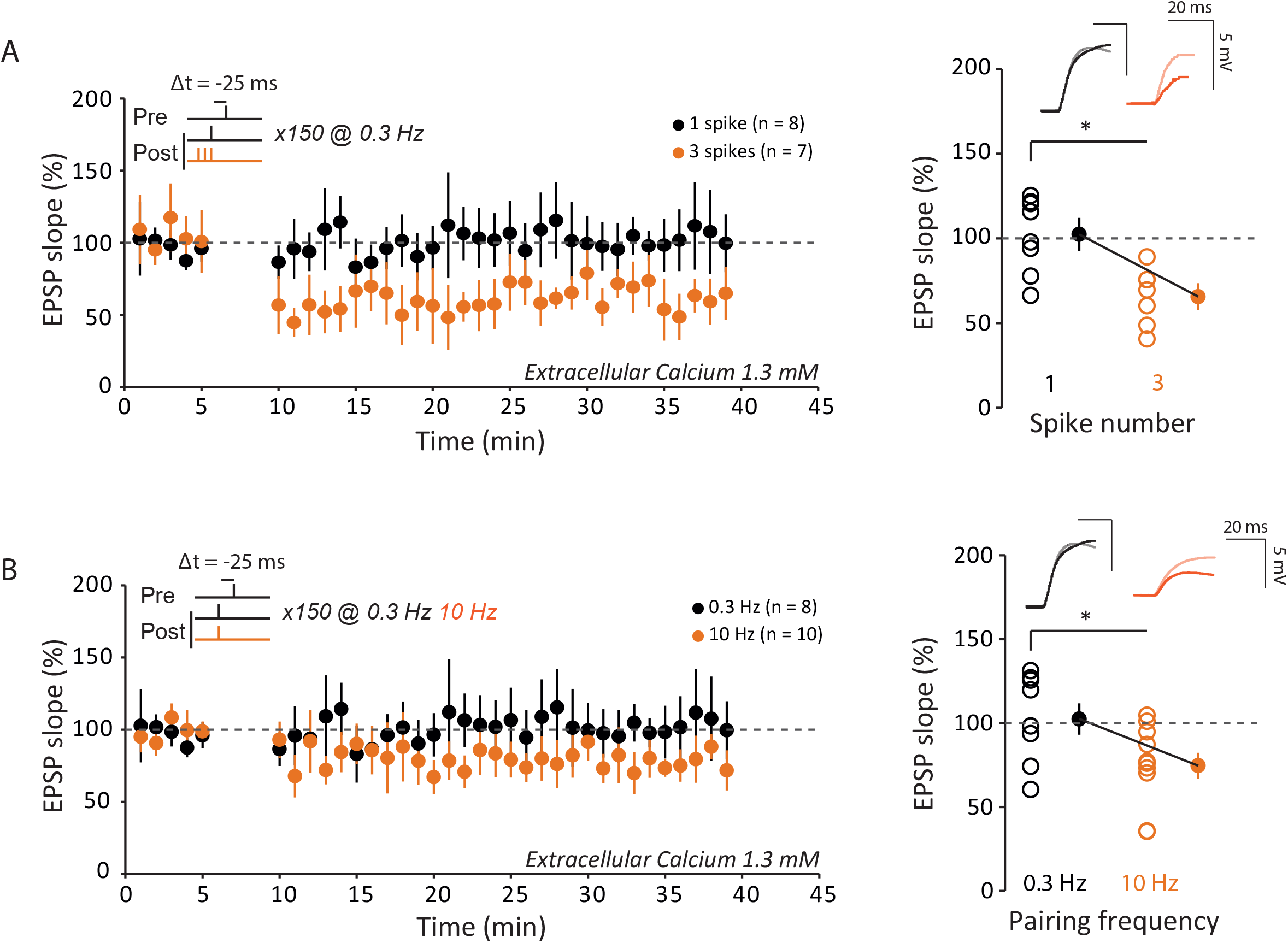
Recovery of t-LTD in 1.3 mM. **(A)** Role of post-synaptic spike number on synaptic changes induced by a post-pre protocol at −25 ms at a pairing frequency of 0.3 Hz in 1.3 mM extracellular calcium. For a single action potential, no change is induced (black). For three action potentials, t-LTD is induced (orange). Thus, increasing the number of post-synaptic action potentials allows t-LTD to recover. **(B)** Effects of stimulation frequency on the induction of synaptic plasticity by a post-pre protocol at −25 ms with a single post-synaptic action potential in 1.3 mM extracellular calcium. For 0.3 Hz, no change is induced (black). For 10 Hz, t-LTD is induced (orange). Thus, increasing pairing frequency allows t-LTD to recover.

### Calcium-based synaptic plasticity model

We built a calcium-based model to quantitatively account for the experimental findings (see Methods for details of the model). Model parameters were fit to the results of the spike-pair protocols at a low pairing-frequency, and the model was then validated by making predictions for the burst and high frequency protocols. Following Graupner and Brunel (2012), our model describes two variables: the intracellular calcium transients in the post-synaptic spine resulting from the pre- and post-synaptic activity, and the dynamics of the synaptic efficacy (or synaptic weight).

Calcium fluxes mediated by NMDAR and VDCC are modeled as discrete jumps following each pre- and post-synaptic spike, respectively. The transients decay with timescale τ_Ca_ and their amplitudes are *C_pre_* and *C_post_* (see Methods). The extra-cellular calcium concentration enters our model only through scaling exponents *a_pre_* and *a_pos_*_t_ of the amplitude parameters *C*_*pre*_ ∝ [*Ca*^2+^]^*apre*^, *C*_*post*_ ∝ [*Ca*^2+^]^*apost*^. These exponents allow our model to account for direct and indirect effects of the extracellular calcium concentration on the calcium transient. Directly, higher concentration leads to increased calcium ion flux upon opening of Ca^2+^ channels. There are also indirect effects including, for example, the dependence of neurotransmitter release probability on [Ca^2+^] ^26^, which in turn affects the postsynaptic transients.

The model is characterized by a second variable, the synaptic weight, that increases at rate *γ_p_* when the calcium transient *c(t)* crosses the potentiation threshold *θ_p_*, and decreases with rate *γ_d_* when *c(t)* crosses the depression threshold *θ_d_*. This is consistent with the notion that high levels of calcium lead to t-LTP, whereas intermediate levels lead to t-LTD ^22, 27, 28^. We assumed that synaptic weight dynamics are graded, i.e., any value of the weight between a lower and upper bound is stable (see Methods).

A hallmark of the biophysics of calcium entry is its non-linearity: when pre- and post-synaptic neurons are co-active, calcium transients are markedly larger than what would be expected due to the sum of independent contributions ^16, 29, 30^. This nonlinearity was characterized in the model by a single parameter, *η*. It is known that calcium transients attributed to nonlinear processes (such as dendritic NMDA spikes) decay on a longer timescale than transients following single spikes ^30, 31^, so the nonlinear term decays with timescale *τ_Ca,NMDA_ > τ_Ca_* (see Methods). Unless noted otherwise, the intracellular calcium transient c(t) is the sum of the above three contributions: pre- and post-synaptic, and the nonlinear term.

### Nonlinear calcium transients are necessary to account for plasticity at multiple extracellular calcium concentrations

A model where calcium dynamics are linear, i.e., pre- and post-synaptic activity contribute to *c(t)* independently is qualitatively consistent with the results of our plasticity experiments: the same relative timing of a pair of pre and postsynaptic spikes may lead to high, intermediate and low levels of transient intracellular calcium at [Ca^2+^] = 3, 1.8, and 1.3 mM; potentially leading to t-LTP, t-LTD and no plasticity, respectively (see Supplementary Figure 2 for illustration of the calcium transients). Furthermore, the linear model was shown to accurately fit plasticity experiments at a single calcium concentration ^23^.

We therefore asked whether such a model could quantitatively fit the data at multiple extracellular concentrations. We find that a linear model does not account for a number of important features of the data (Supplementary Figure 3). In particular, the inferred decay timescale of calcium is long (*τ_Ca_* = 76 ms, see Tables 1, 2 for full list of model parameter values), so at 3 mM (Supplementary Figure 3A, top) the model predicts significant changes to synaptic weights even when the pre- and post-synaptic spikes are separated by more than 100 ms, inconsistent with our data as well as other experiments ^6, 24^. Furthermore, the long decay timescale and the small difference between the depression and potentiation thresholds implies that under many protocols with multiple post-synaptic spikes, or at high pairing frequencies, the synaptic weight saturates to its upper bound. In the experiment, the same protocols resulted in t-LTD, leading to large inaccuracies in the predictions (Supplementary Figure 3B, C). This model also failed to reproduce the dependence of calcium entry on the extra-cellular calcium concentration measured in imaging experiments (Supplementary Figure 4) unless non-realistic scaling exponents (*a_pre_, a_post_*) were used (see Methods).

**Table 1.** Parameters and resulting fitting and prediction errors for models chosen based on best combined error for pair and burst plasticity protocols.

|  |  | Model (imaging constraints) |  |  |  |
| --- | --- | --- | --- | --- | --- |
| Param. | Meaning | nonlinear<br>(none) | nonlinear<br>(2 s.d.) | nonlinear<br>(1 s.d.) | linear<br>(none) |
| $C_{pre}$ | Pre-synaptic amplitude | 0.105 | 0.135 | 0.755 | 0.622 |
| $C_{post}$ | Post-synaptic amplitude | 0.127 | 0.570 | 0.189 | 0.340 |
| $a_{pre}$ | Pre-syn. $Ca^{2+}$ exponent | 0.594 | 0.859 | 0.111 | 0 |
| $a_{post}$ | Post-syn. $Ca^{2+}$ exponent | 1.538 | 0.499 | 1.294 | 0.966 |
| $\tau_{Ca}$ (ms) | $Ca^{2+}$ timescale | 96.040 | 18.185 | 33.961 | 75.753 |
| $D$ (ms) | Pre-synaptic delay | 15.473 | 0.942 | 8.668 | 7.412 |
| $\theta_p$ | Potentiation threshold | 5.834 | 3.002 | 1.173 | 1.326 |
| $\gamma_d$ | Depression rate | 0.122 | 1.212 | 0.388 | 0.047 |
| $\gamma_p$ | Potentiation rate | 0.944 | 1.052 | 1.998 | 0.332 |
| $w_{min}$ | Minimum synaptic weight | 0.829 | 0.840 | 0.833 | 0.781 |
| $w_{max}$ | Maximum synaptic weight | 1.411 | 2.241 | 1.344 | 1.394 |
| $\tau_{Ca,NMDA}$ (ms) | Nonlin. $Ca^{2+}$ timescale | 241.521 | 128.923 | 162.420 | N/A |
| $\eta$ ($ms^{-1}$) | Nonlinearity parameter | 410.352 | 414.466 | 0.00436 | N/A |
| $\epsilon_{pair}$ | Pre-post pair error | 0.203 | 0.227 | 0.229 | 0.196 |
| $\epsilon_{burst}$ | Post-synaptic burst error | 0.317 | 0.326 | 0.320 | 0.414 |
| $\epsilon_{total}$ | Total low freq. plasticity error (pair & burst) | 0.267 | 0.281 | 0.279 | 0.324 |
| $\epsilon_{freq}$ | High freq. plasticity error | 0.405 | 0.344 | 0.424 | 0.370 |
| $\epsilon_{imaging}$ | Imaging error | 1.219 | 0.971 | 0.877 | 0.872 |
| Corresponding figure |  | S. Fig. 5 | Fig. 7<br>Fig. 8<br>S. Fig. 2<br>S. Fig. 3 | S. Fig. 5 | S. Fig. 2<br>S. Fig. 3 |

**Table 2.** Parameters and resulting fitting and prediction errors for models chosen based on best error for pair plasticity protocols.

|  |  | Model (imaging constraints) |  |  |  |
| --- | --- | --- | --- | --- | --- |
| Param. | Meaning | nonlinear<br>(none) | nonlinear<br>(2 s.d.) | nonlinear<br>(1 s.d.) | linear<br>(none) |
| $C_{pre}$ | Pre-synaptic amplitude | 0.0108 | 0.446 | 0.558 | 0.380 |
| $C_{post}$ | Post-synaptic amplitude | 0.401 | 0.141 | 0.138 | 0.554 |
| $a_{pre}$ | Pre-syn. $Ca^{2+}$ exponent | 2.288 | 0.681 | 0.426 | 0.234 |
| $a_{post}$ | Post-syn. $Ca^{2+}$ exponent | 0.643 | 1.566 | 1.560 | 0.319 |
| $\tau_{Ca}$ (ms) | $Ca^{2+}$ timescale | 70.129 | 17.946 | 41.087 | 191.513 |
| $D$ (ms) | Pre-synaptic delay | 20.951 | 7.169 | 23.675 | 6.936 |
| $\theta_p$ | Potentiation threshold | 5.633 | 3.816 | 1.145 | 1.174 |
| $\gamma_d$ | Depression rate | 1.083 | 1.133 | 1.954 | 0.239 |
| $\gamma_p$ | Potentiation rate | 0.966 | 0.439 | 0.660 | 2 |
| $w_{min}$ | Minimum synaptic weight | 0.793 | 0.816 | 0.778 | 0.776 |
| $w_{max}$ | Maximum synaptic weight | 2.736 | 3 | 3 | 1.392 |
| $\tau_{Ca,NMDA}$ (ms) | Nonlin. $Ca^{2+}$ timescale | 92.842 | 149.217 | 172.758 | N/A |
| $\eta$ ($ms^{-1}$) | Nonlinearity parameter | 342.891 | 434.382 | 0.00619 | N/A |
| $\epsilon_{pair}$ | Pre-post pair error | 0.199 | 0.218 | 0.229 | 0.194 |
| $\epsilon_{burst}$ | Post-synaptic burst error | 0.358 | 0.344 | 0.349 | 0.505 |
| $\epsilon_{total}$ | Total low freq. plasticity error | 0.290 | 0.288 | 0.295 | 0.383 |
| $\epsilon_{freq}$ | High freq. plasticity error | 0.445 | 0.299 | 0.417 | 0.414 |
| $\epsilon_{imaging}$ | Imaging error | 1.349 | 0.929 | 0.887 | 1.005 |
| Corresponding figure |  | S. Fig. 6 | S. Fig. 6 | S. Fig. 6 | N/A |

### A nonlinear calcium based model generates accurate predictions for post-synaptic burst and high-frequency induction protocols

We therefore focus henceforth on models that include a nonlinear contribution to calcium transients (Figure 7). Here, the inferred decay timescale of transients originating from a single neuron is short (*τ_Ca_* = 18 ms, see Table 1), which implies that pre- and post-synaptic spikes must be relatively close in time to elicit a transient that exceeds the potentiation and depression threshold. Thus, significant changes to synaptic weights in STDP protocols are restricted to *Δt* between −50 and 50 ms (for pre-post pairs, Figure 8A). The model correctly captures the dependence of plasticity on the number of spikes in a post-synaptic burst, and on its timing (Figure 8B). Specifically, our model correctly predicts that at [Ca^2+^] = 1.8 mM, increasing the number of post-synaptic spikes that follow a pre-synaptic spike from one to four leads to a change in the direction of plasticity from t-LTD to t-LTP; and that a single pre-synaptic spike followed by a burst of three post-synaptic spikes leads to t-LTP, while the reverse order of the same stimulation pattern leads to t-LTD.

**Figure 7.**
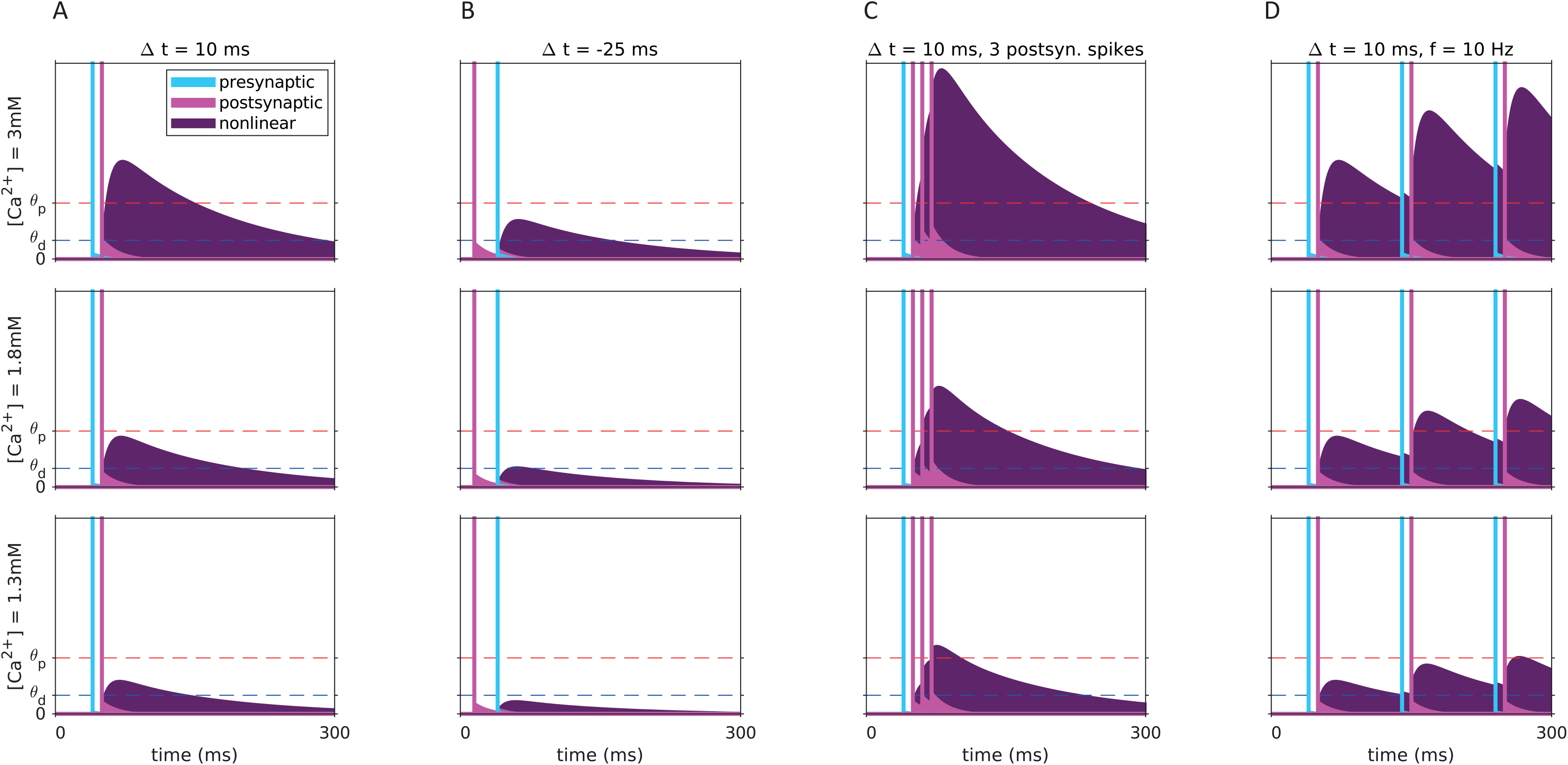
Dependence of calcium transients of nonlinear plasticity model on the stimulation protocol and on the extracellular calcium concentration. Example model calcium transients resulting from a single pre-post pairing at Δt = 10 ms (A), Δt = −25 ms (B), a pre-synaptic spike paired with a burst of three post-synaptic spikes (C) and a pre-post pair repeated at a frequency of 10 Hz (D). Transients for all stimulation protocols are shown for the three extracellular calcium concentrations used in the experiment (rows). The linear pre-and post-synaptic contributions are shown in cyan and magenta, respectively, together with the spike times indicated by the vertical lines. The nonlinear contribution is shown in purple. The transients depend strongly on the relative timing of pre- and post-synaptic activity, primarily due to the model’s nonlinearity. The long decay timescale of the nonlinear term implies that calcium returns to baseline slow and thus may accumulate for induction protocols with high pairing frequencies.

**Figure 8.**
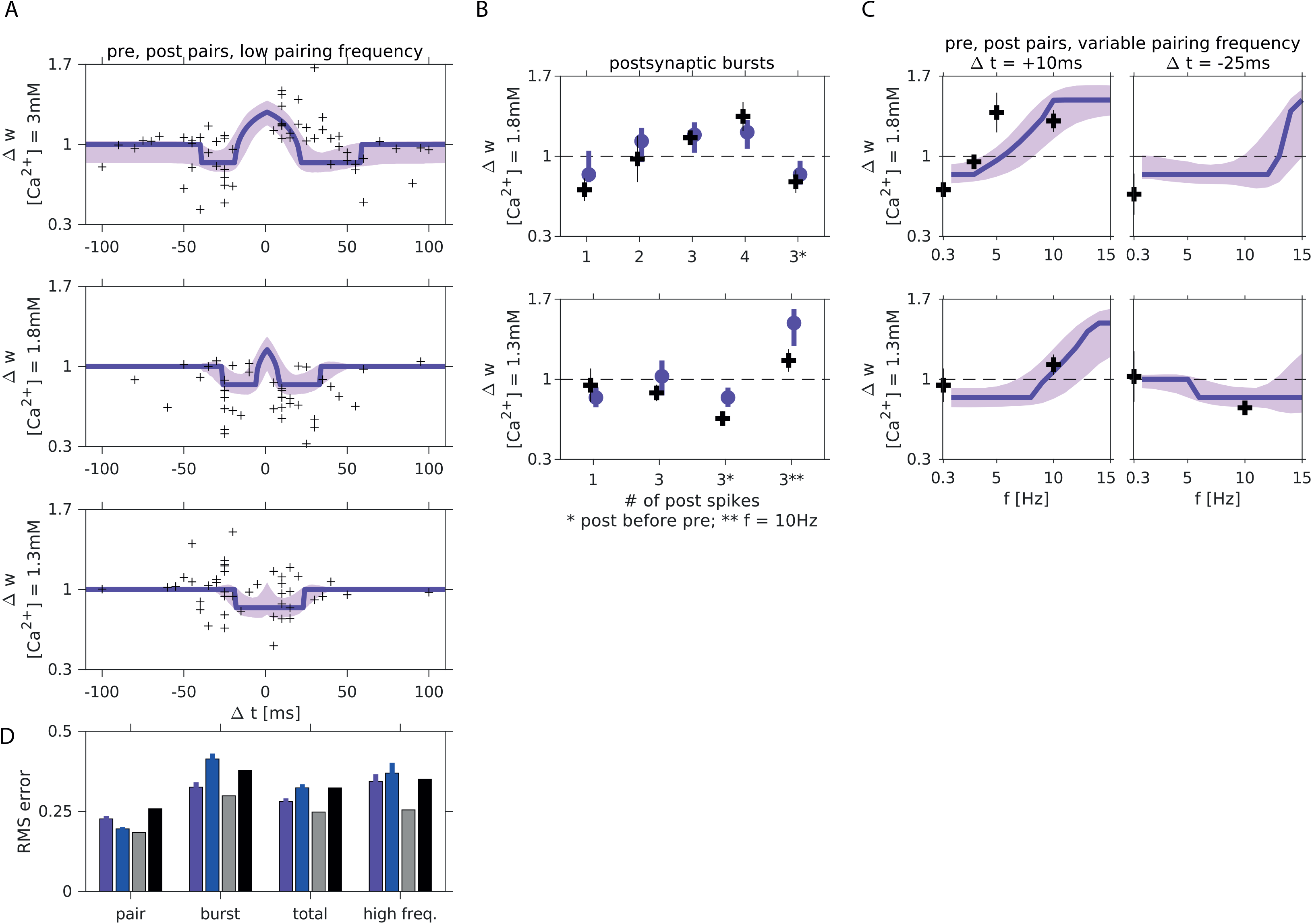
A nonlinear calcium-based plasticity model fits well plasticity results at multiple concentrations and using multiple induction protocols. **(A)** Model fit to the STDP data at [Ca^2+^] = 3, 1.8, 1.3 mM (purple). Each measured synapse is represented by a black cross. Shaded area indicates the standard deviation around the mean obtained by generating predictions using parameter sets in the neighborhood of the best fitting model (see Methods). **(B)** Experimental results and model predictions for protocols with post-synaptic burst stimuli. At [Ca^2+^] = 1.8 mM (top), the model captures the change in sign of plasticity as a function of the number of spikes in the post-synaptic burst, and the reverse in sign when a post-synaptic burst of three spikes precedes a pre-synaptic spike. At [Ca^2+^] = 1.3 mM (bottom), LTD is restored at a low pairing-frequency by a burst of three post-synaptic spikes (regardless if they arrive closely before or after the pre-synaptic spike), compared to a spike-pair protocol where no significant change was observed, which is captured by the model. **(C)** Experimental results and model predictions for protocols at variable pairing frequencies. For spike pair protocols with Δt = 10 ms, the resulting plasticity depends strongly on the pairing frequency. At [Ca^2+^] = 1.8 mM (top) the model accurately predicts the frequency at which plasticity switches sign. At [Ca^2+^] = 1.3 mM the model correctly predicts the change in sign of plasticity when pre- and post-synaptic spikes are paired at 10 Hz with positive/negative relative timing. **(D)** Root mean square (RMS) errors when the model predictions are compared to the spike-pair data, the bust data (only at a low pairing frequency of 0.3 Hz), and measurements of plasticity using high-frequency pairing protocols. Total error refers to the combined error for spike-pair and burst stimuli at (a low pairing frequency). Note that the model was fit using the spike-pair data. Among all fitted parameter sets, this figure shows results for the model with lowest combined spike-pair and burst protocols. Results from high-frequency protocols (panel C and parts of panel B) were not used in fitting or model selection. See Supplementary Figure SI_M_PAIR for fit and predictions of the model with lowest RMS error on spike-pair data. Error bars for model results, predictions and errors (panel D) are equal to the standard deviation of the RMS error computed 200 times on subsets (80% of the data chosen randomly) of data points. Gray bars show estimates of the variability of the data, and black bars show the errors made by a null model (see Table 4).

Furthermore, owing to the long decay timescale associated with the nonlinear term (*τ_Ca,NMDA_* = 129 ms), the model correctly predicts the frequency dependence of plasticity (Figure 8C, these data points were not used in fitting; see Methods). At [Ca^2+^] = 1.8 mM it correctly predicts the frequency at which a protocol with pair of spikes with *Δt* = 10 ms switches from t-LTD to t-LTP (Figure 8C, top), and at [Ca^2+^] = 1.3 mM it correctly predicts the difference in sign of plasticity between protocols with *Δt* = 10 ms and −25 ms (Figure 8C, bottom).

The inferred model suggests that changes to synaptic efficacy induced by the STDP protocol (controlled by the rates *γp*, *γd*) are large relative to the range of possible efficacy values (*w_max_ – w_min_*). Such saturation leads to an STDP curve with a square shape (Figure 7), and suggests that a smaller number of pairings could be sufficient to induce saturation, especially to the lower bound of a synapse’s efficacy. This is consistent with experimental tests of the possible discrete nature of plasticity in these synapses ^32, 33^. This abrupt switch between t-LTD and t-LTP is nevertheless consistent with the typical smooth shape of STDP curves, if heterogeneities are included (Figure 2 B, C, D). To show that, we generated STDP curves by randomly varying the model parameters around the best fitting set (see Methods) and plotted one standard deviation above and below the average STDP curve, yielding a curve similar to the experimental one (Figure 8A, light purple shaded region). This reflects the fact that each point in our dataset corresponds to a measurement of a different synapse, and suggests that the parameters found by our fitting procedure are not fine-tuned.

Importantly, model parameters were fit to the STDP results (i.e., plasticity resulting from pre-post pairs at different *Δt*), while satisfying constraints imposing consistency with the dependence of the total calcium entering the cell on the extracellular concentration (Supplementary Figure 4 B, C). The mathematical form of these constraints was derived analytically based on the model structure, and the values used in the inequalities were based on our imaging experiments (see Methods). Imposing such constraints on the model parameters led to a small increase of the fitting error (Supplementary Figures 5, 6; Tables 1, 2). However, interestingly, constraints based on imaging experiments yield models with more realistic windows for induction of STDP (Supplementary Figures 5A, 6A, top) and more accurate predictions for the frequency dependence of plasticity (Figure 8D, Supplementary Figures 5D, 6D).

### Residual LTD at [Ca^2+^] = 1.3 mM

The STDP curve at the lowest concentration we measured showed non-statistically significant LTD at positive relative timing (Figure 2E, see results for [Ca^2+^] = 1.3 mM, *Δt* = +10 ms). The models we inferred from the data show a small window for the induction of LTD at 1.3 mM (Figure 8A, bottom). We used the model to predict the maximum concentration for which a pair of spikes at timing *Δt* = +10 ms will not lead to transient calcium crossing the depression threshold, and found that to be [Ca^2+^] = 1.13 mM. We note that the window for induction of LTD exhibited by our model is consistent with the data, when the variability is taken into account (Figure 8A, bottom). Models constrained to exhibit no LTD at [Ca^2+^] = 1.3 mM fit poorly to the STDP curves at higher concentrations (not shown).

## Discussion

Our study reveals that the STDP curve at the Schaffer collateral-to-CA1 pyramidal cell synapses measured in external physiological calcium (1.3-1.8 mM) and in the presence of normal synaptic inhibition does not follow the classical asymmetric shape reported at non physiological calcium (2.5-3 mM). Rather, we show that no significant plasticity occurs in the lower range of the physiological calcium range, while only depression occurs in the upper range, when single EPSPs and post-synaptic spikes are paired. Thus, there is a complete lack of potentiation at external Ca^2+^ concentration ranging between 1.3 and 1.8 mM. We then find that t-LTP can be restored in physiological Ca^2+^ either by increasing the number of post-synaptic spikes (from 1 to 3-4) or by increasing the pairing frequency (from 0.3 to 10 Hz). Likewise, t-LTD can be restored at 1.3 mM by increasing the number of post-synaptic spikes during the pairing from 1 to 3.

### t-LTP and physiological external calcium

We report here that t-LTP could not be induced by positive correlation between single pre- and post-synaptic spikes in physiological external calcium. Rather, t-LTD and no plasticity were respectively induced at 1.8 and 1.3 mM calcium. It is noteworthy that for positive delay (+5/+25 ms), the synaptic plasticity goes from no change at the lower limit of calcium range (1.3 mM) through a depression phase at the upper limit of calcium range (1.8 mM) before reaching the potentiation level for non-physiological, high calcium (2.5 and 3 mM). Such biphasic behavior was predicted by calcium-based theoretical studies of synaptic plasticity ^22, 23, 27^.

t-LTP at positive delay could be restored in physiological calcium by either increasing the number of postsynaptic spikes from 1 to 3 or 4 (at 1.8 mM but not at 1.3 mM calcium) or by increasing the pairing frequency from 0.3 to 10 Hz (at 1.3 or 1.8 mM calcium). Such recovery of t-LTP had been already reported in previous studies ^18, 25^, but the data reported in these studies were obtained in non-physiological calcium and in the absence of synaptic inhibition. This finding suggests that the minimal paradigm for inducing robust t-LTP *in vivo* would be to pair EPSPs associated with spike bursting at theta frequency. In fact, this type of pairing has been shown to induce synaptic potentiation of thalamo-cortical EPSPs in the cat visual cortex *in vivo* ^34^.

### t-LTD and physiological external calcium

At 3 mM calcium, t-LTD was induced by negative delays (−5/−50 ms) and for positive delays (+40/+60 ms). This second t-LTD window has been predicted by calcium-based models ^22, 23, 35^ and had been reported previously in a few experimental studies in which calcium transients were enhanced by spike broadening ^8, 13^ or by multiple spike-pairing at 5 Hz ^18^. t-LTD was also observed for both negative and positive delays at 1.8 mM, but no significant t-LTD was observed for negative delays (−5/−25 ms) at 1.3 mM and 1.5 mM calcium. Nevertheless, t-LTD could be restored at 1.3 mM calcium when the number of postsynaptic spikes was increased from 1 to 3.

### Comparison of experimental data and calcium-based models

The successive transitions from a STDP curve with both t-LTD and t-LTP first to a curve with t-LTD only, and then to a curve with no plasticity, are generic predictions of calcium-based models in which potentiation occurs at high calcium concentrations, while depression occurs at intermediate calcium concentrations ^22, 23^. Furthermore, these models also generically predict that increasing frequency, or number of spikes in a burst, will lead eventually to t-LTP ^22, 23^. Here, we have a made a major step beyond these generic predictions, by fitting quantitatively experimental data at various calcium concentrations with variants of calcium - based synaptic plasticity models (linear and non-linear). The data reported here measured at multiple concentrations allowed us to show that non-linear summation of transient calcium contributions is necessary to reproduce quantitatively the experimental findings. We chose the nonlinear interaction term to be quadratic for simplicity. This nonlinearity describes in a schematic fashion the biophysical properties of the NMDA receptor, that needs a coincidence of both presynaptic (for glutamate binding) and postsynaptic (for magnesium block relief) activity in order to maximize opening probability. It has been directly observed in calcium imaging experiments (e.g. ^16, 29^), and is necessary to explain our own imaging experiments, comparing calcium transients following positive and negative timings (Figure 3, Supplementary Figure 1, 4).

Furthermore, we found the non-linear term should have a decay timescale of approximately 100 ms (longer than that of transients due to pre- or postsynaptic activity alone, ∼20 ms) for the model to make accurate predictions for high pairing-frequency protocols. These conclusions are consistent with the work of Brandalise et al. (2016) where it was shown that dendritic NMDA spikes following coincident stimulation of pre and post synaptic activity lead to calcium transients that decay slowly, and that blocking these NMDA spikes abolishes t-LTP measured in the control protocol. Note however that these experiments were carried out in recurrent CA3 synapses and not in CA3 to CA1 synapses.

These results call for a radical reevaluation of classic ‘STDP rules’ (and of the experimental conditions under which they are measured), in which pairs of single pre and postsynaptic spikes modify synaptic efficacy; rather, they show that such pairs (even if repeated many times at low frequency) are not likely to elicit any changes of synaptic strength in physiological conditions. They indicate that plasticity can only be triggered when bursts of spikes are used, or neurons fire at sufficiently high frequency, and invalidate popular phenomenological models of STDP in which plasticity is triggered by pairs of single pre and post-synaptic spikes. Qualitatively similar findings have been obtained in synaptic plasticity experiments in granule cell to Purkinje cell synapses, where bursts of pre-synaptic spikes are required to induce both t-LTD ^35^ and t-LTP ^36^, and in area RA of songbirds ^37^. In cortical slices, it has been shown that pairing single spikes at low frequencies only induces LTD but not LTP _17_.

Beyond spiking activity, the recent work of Bittner et al. (2017) serves as another example of how LTP can be induced reliably in physiological conditions: these authors have shown that a plateau potential in a CA1 neuron can lead to potentiation (and, in fact, creation of a new place field) even when the preceding pre-synaptic activity occurs up to a second before it ^38^. The large calcium influx during the plateau potential suggests that it might be possible to construct a biophysical, calcium-based model that will parsimoniously describe plasticity due to both spiking activity and plateau potentials in the hippocampus.

## Methods

### Acute slices of rat hippocampus

Hippocampal slices were obtained from 14- to 20-day-old rats according to institutional guidelines (Directive 86/609/EEC and French National Research Council) and approved by the local health authority (# D1305508, Préfecture des Bouches-du-Rhône, Marseille). Rats were deeply anaesthetized with chloral hydrate (intraperitoneal 400mg/kg) and killed by decapitation. Slices (350 µm) were cut in a N-methyl-D-glucamine (NMDG)-solution (in mM: 92 NMDG, 2.5 KCl, 1.2 NaH_2_PO_4_, 30 NaHCO_3_, 20 HEPES, 25 glucose, 5 sodium ascorbate, 2 thiourea, 3 sodium pyruvate, 10 MgCl_2_, 0.5 CaCl_2_) on a vibratome (Leica VT1200S) and were transferred at 32°C in NMDG-solution for 20 min before resting for 1 h at room temperature in oxygenated (95% O_2_/5% CO_2_) artificial cerebro-spinal fluid (ACSF; in mM: 125 NaCl, 2.5 KCl, 0.8 NaH_2_PO_4_, 26 NaHCO_3_, x CaCl_2_, x MgCl_2_, and 10 D-glucose). The ratio of CaCl_2_ and MgCl_2_ concentration was maintained constant to stabilize presynaptic release probability ^39, 40^ and external Ca^2+^ concentration was set to either 3 mM (Mg^2+^: 2 mM), 1.8 mM (Mg^2+^: 1.2 mM) or 1.3 mM (Mg^2+^: 0.9 mM). For recording, each slice was transferred to a temperature-controlled (30°C) chamber with oxygenated ACSF.

### Organotypic slices of rat hippocampus

Postnatal day 5-7 Wistar rats were deeply anesthetized by intraperitoneal injection of chloral hydrate, the brain was quickly removed, and each hippocampus was individually dissected. Hippocampal slices (350µM) were placed on 20 mm latex membranes (Millicell) inserted into 35 mm Petri dishes containing 1 ml of culture medium and maintained for up to 21 days in an incubator at 34°C 95% O_2_-5% CO_2_. The culture medium contained (in ml) 25 MEM, 12.5 HBSS, 12.5 Horse serum, 0.5 penicillin/streptomycin, 0.8 glucose (1M), 0.1 ascorbic acid (1mg/ml), 0.4 Hepes (1M), 0.5 B27 and 8.95 sterile H_2_O. To avoid glial proliferation, 5 µM Ara-C was added to the medium at 3 days in vitro (DIV) for one night. Pyramidal cells from the CA1 arena were recorded at the age of 8-10 DIV.

### Electrophysiology

Neurons were identified with an Olympus BX 50WI microscope using Differential Interference Contrast (DIC) x60 optics. Whole-cell recordings were made from CA1 pyramidal neurons, electrodes were filled with a solution containing the following (in mM): 120 K-gluconate, 20 KCl, 10 HEPES, 2 MgCl_2_6H_2_O, and 2 Na_2_ATP. Stimulating pipettes filled with extracellular saline were placed in the stratum radiatum (Schaffer collaterals). In control and test conditions, Excitatory Post-Synaptic Potentials (EPSPs) were elicited at 0.1 Hz by a digital stimulator that fed a stimulation isolator unit (A385, World Precision Instruments). Access resistance was monitored throughout the recording and only experiments with stable resistance were kept (changes<20%).

### Acquisition and data analysis

Recordings were obtained using a Multiclamp 700B (Molecular Devices) amplifier and pClamp10.4 software. Data were sampled at 10 kHz, filtered at 3 kHz, and digitized by a Digidata 1440A (Molecular Devices). All data analyses were performed with custom written software in Igor Pro 6 (Wavemetrics). EPSP slope was measured as an index of synaptic strength. Pooled data are presented as mean ± SEM. Statistical comparisons were made using Wilcoxon or Mann-Whitney test as appropriate with Sigma Plot software. Data were considered as significant when P<0.05.

### STDP induction protocols

After obtaining a stable EPSP baseline for a period of 10-15 minutes, single EPSPs were paired with a STDP protocol was applied on the input with 100 repetitions for LTP and 150 repetitions for LTD. The number of post-synaptic spikes and the pairing frequencies were adjusted depending on the experiment. The post-synaptic spikes(s) were evoked by a brief somatic current pulse. EPSP slopes were monitored at least for 20 minutes after each pairing episode. Values of synaptic change were measured in the time window between 15 and 25 minutes after pairing.

### Calcium imaging

CA1 pyramidal neurons from organotypic slice cultures of hippocampus were imaged with a LSM-710 Zeiss confocal microscope. For imaging calcium in the spine, 50 µM Alexa-594 and 250 µM Fluo-4 (Invitrogen) were added to the pipette solution. Alexa-594 fluorescence was used to reveal neuronal morphology, whereas fluorescence signals emitted by Fluo-4 were used for calcium imaging. Laser sources for fluorescence excitation were set at 488 nm for Fluo-4 and 543 nm for Alexa-594. Emitted fluorescence was collected between 500 and 580 nm for Fluo-4 and between 620 and 750 nm for Alexa-594. After whole-cell access, the dyes were allowed to diffuse for at least 10 minutes before acquisition of fluorescence signals. EPSP were evoked by a digital stimulator (A385, World Precision Instruments) and action potentials were elicited by a pulse of depolarizing current. Electrophysiological signals were synchronized with calcium imaging in line-scan mode. Acquired Fluo-4 signals were converted to ΔG/R values and peak amplitudes were measured with a custom-made software (LabView, National Instruments).

### Synaptic plasticity model

#### Calcium transients

Calcium transients in the model depend on the pre- and post-synaptic spiking activity, and indirectly on the extracellular calcium concentration (through scaling of the amplitude parameters with [Ca^2+^]). The pre- and post-synaptic spike trains are denoted by *s*_*pre*_ (*t*) = ∑_*i*_ *δ*(*t* − *t*_*i*,*pre*_) and *s*_*post*_ (*t*) = ∑_*i*_ *δ*(*t* − *t*_*i*,*post*_), respectively. The temporal evolution of the pre- and post-synaptically induced calcium transients is given by:

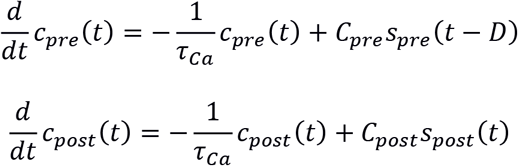

where *C_pre_* and *C_post_* are parameters describing the magnitude of jumps in calcium following a pre- and a post-synaptic spike, D is a pre-synaptic delay parameter and *τ_Ca_* is the decay timescale of calcium transients.

The equation for the time evolution of the nonlinear contribution *c_NL_(t)* is

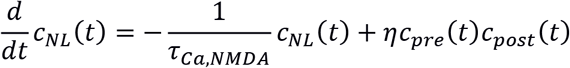

where *τ_Ca,NMDA_* is the decay timescale of the nonlinear transients and *η* is a parameter describing the strength of the nonlinearity. The total transient *c(t)* is then

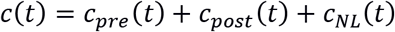

The amplitude parameters *C_pre_*, *C_post_* depend on the extracellular calcium concentration through the scaling exponents *a_pre_*, *a_post_*:

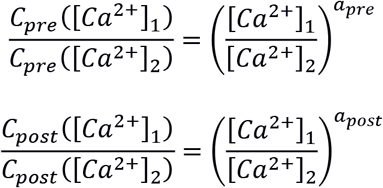

In addition to the full nonlinear model described by the equations above, we considered linear models where the calcium transient is a sum over pre- and post-synaptic contributions only, i.e., *c(t) = c_pre_(t) + c_post_(t)* (equivalently, one can set *η = 0*).

We also considered a model in which there is no linear post-synaptic contribution. The equations for the time evolution of calcium transients in that case are the same as above, with *c(t) = c_pre_(t) + c_NL_(t)* (note that the post-synaptic transient must still be computed to then compute the nonlinear one).

#### Synaptic weight dynamics

In our model, synaptic weight dynamics are graded, i.e., any synaptic weight value between *w_min_* and *w_max_* is stable if there is no crossing of the depression (and potentiation) threshold. The weight variable w is restricted to this range using soft thresholds, so the equation for the time evolution of *w* is

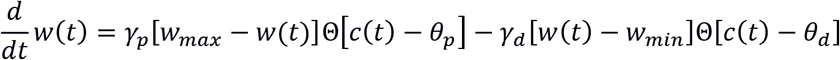

Here, *ϴ* is the heaviside step function; *γ_p_* and *γ_d_* are the potentiation and depression rates, respectively; and *θ_p_* and *θ_d_* are the potentiation and depression thresholds above which the synaptic weight variable increases or decreases.

#### Using the imaging experiments to constrain model parameters

In the imaging experiments, we measured the calcium fluorescence in postsynaptic spine under different stimulation protocols and in preparations with differing calcium concentrations. We used the results of these experiments to constrain the parameters of the calcium based plasticity model. Specifically, we computed the area under the ΔG/F curve of resulting from protocols with one pre- and 1, 2, 3 or 4 post-synaptic spikes (Figures 3, 4), measured at different extracellular calcium concentrations. This is thought to be proportional to the total calcium entry following a given stimulus at a given concentration. The quantity that was related to data was then the ratio of total calcium entry for a two post-synaptic spike stimulus relative to a protocol with a single post-synaptic spike (denoted *r_2_*). At [Ca^2+^] = 1.3 mM we have *r_2_* = 1.05 ± 0.26, while at [Ca^2+^] = 3 mM we have *r_2_* = 1.88 ± 0.42 (mean ± standard deviation). We also considered measurements of calcium transients following pre-posts pairs of spikes with positive and negative timing (*Δt* = ±20 ms). Specifically, we computed the ratio *r_±_* of the corresponding ΔG/F curves and found *r_±_* = 1.51 ± 0.63 for [Ca^2+^] = 1.3 mM and 1.88 ± 0.43 for [Ca^2+^] = 3 mM.

#### Linear case

When calcium transients are linear in the pre and post-synaptic activity, the model parameters can be constrained to reproduce the values of *r_2_* measured in the experiment. These constraints are (see **Supplementary Mathematical Note** for derivation): *C_post_/C_pre_* = 0.011 and *a_post_ - a_pre_* = 5.93. Assuming *a_post_* and *a_pre_* are non-negative (i.e., calcium amplitudes do not decrease with the extracellular concentration) implies *a_post_* ≥ 5.93. We think that such a scaling exponent is not biophysically realistic, so we interpret the imaging experiments as additional evidence that nonlinear calcium transients must be taken into account in the plasticity model.

#### Nonlinear case

We expressed *r_2_* using the parameters of the nonlinear model (see Supplementary Mathematical Note for derivation and formula), and fit model parameters subject to inequalities guaranteeing that the model’s *r_2_* is within one or two standard deviations of its mean, computed from the experimental results. The model fits, predictions and errors are shown in Figure 8 and Supplementary Figures 5, 6.

#### Constraints on parameters from qualitative features of the model and the STDP curve

For the transient amplitude parameters *C_pre_* and *C_post_* the constraints we used ensure that a single spike by a pre- or post-synaptic neuron cannot lead to a calcium transient that exceeds the depression threshold at the highest calcium concentration we used in our experiments ([Ca^2+^] = 3.0 mM).

This constraint can be relaxed if the model is fitted to data with a single calcium concentration. One can then ensure that a pair of well separated pre- and post-synaptic spikes do not lead to changes in the synapse by requiring that the product of the time spent above threshold (*T_p_*, *T_d_*) and the corresponding rate (*γ_p_*, *γ_d_*) are equal. If transients induced by single spikes can cross threshold, this requires tuning either *γ_p_* or *γ_d_* such that indeed *γ_d_ T_d_ = γ_p_ T_p_* ^23^.

This scheme cannot be used when one is interested in fitting the model for multiple extracellular calcium concentrations. The reason is that if one fixes, say, *γ_d_* such that potentiation and depression balance each other for a pair of spikes at one concentration, the same value will lead to imbalance at a different concentration.

We assumed that the calcium transients following a single pre- and post-synaptic spike decay on the same timescale, *τ_Ca_*, while the time-scale associated with the nonlinear term, *τ_Ca,NMDA_*, is longer. This assumption can be justified, at least qualitatively, by inspection of the STDP curve we measured at [Ca^2+^] = 3.0 mM and the fact that increasing the pairing-frequency above a few Hz leads to strong changes in the resulting plasticity.

From the shape of *Δw* as a function of *Δt* at high concentration we conclude that the calcium transients in the model must carry information about the relative timing of pre- and post-synaptic spikes at a resolution of approximately 20 ms. In other words, the fact that changing the relative timing of a pair of spikes by 20 ms leads to significant changes to the resulting plasticity implies that the calcium transients in the model must preserve the timing information on that timescale.

On the other hand, the impact that increasing the pairing-frequency has on the resulting plasticity implies that the model must consist of at least one transient that decays on a timescale significantly longer than 20 ms.

We further inspect the shape of the STDP curve measured at [Ca^2+^] = 3.0 mM. Especially noteworthy is the second LTD window at positive *Δt*. The width of that window is comparable with the first LTD window at negative *Δt*. Moreover, consider reflecting the data about a vertical axis at the center of the LTP window, *Δt* ∼ 20 ms. The resulting data points give an STDP curve that overlaps the original curve, suggesting a degree of symmetry played by the pre-synaptic neuron (including the delay D) and the post-synaptic neuron. This symmetry (which is also respected by the STDP curves measured at [Ca^2+^] = 1.3, 1.8 mM) implies that one cannot associate the pre-synaptic transients with a decay timescale much shorter than that of post-synaptic transients, or vice versa.

Therefore, assuming there are no intermediate timescales between *τ_Ca_* and *τ_Ca,NMDA_* that need to be explicitly introduced to the model, the only way to associate the timescales with transients that respects this symmetry is to let the pre- and post-synaptic transients decay with timescale *τ_Ca_*, and let the nonlinear transient decay with timescale *τ_Ca,NMDA_*. We note that this qualitative argument yields a model that is in good agreement with the known biophysics of calcium entry in hippocampal neurons.

#### Numerical optimization

A detailed description of the procedure and formulas we used to fit the different models is included in the supplementary mathematical note. Table 3 includes the range of values within which we optimized the model parameters, and the values we used (for parameters that were fixed).

**Table 3.** Allowed ranges for parameters in numerical optimization procedure.

| Param. | Meaning | Minimum | Maximum | Comments |
| --- | --- | --- | --- | --- |
| $C_{pre}$ | Pre-synaptic amplitude | 0.01 | 1 | We set additional constraints so that single neuron (pre/post) transients did not cross $\theta_d$ at the highest concentration $[Ca^{2+}] = 3$ mM. |
| $C_{post}$ | Post-synaptic amplitude | 0.01 | 1 | |
| $a_{pre}$ | Pre-syn. $Ca^{2+}$ exponent | 0 | 3 | |
| $a_{post}$ | Post-syn. $Ca^{2+}$ exponent | 0 | 3 | |
| $\tau_{Ca}$ (ms) | $Ca^{2+}$ timescale | 0 | 100, 250 | We used the higher upper bound for the linear model. |
| $D$ (ms) | Pre-synaptic delay | 0 | 40 | |
| $\theta_p$ | Potentiation threshold | 1 ($=\theta_d$) | 10 | The depression threshold was fixed to 1, and transient calcium amplitudes were measured relative to it. |
| $\gamma_d$ | Depression rate | 0.0001 | 2 | We tried increasing the upper bound for $\gamma_d$ leading in some cases to slightly better fitting errors (see Table 2), but worse predictions. |
| $\gamma_p$ | Potentiation rate | 0.0001 | 2 | |
| $w_{min}$ | Minimum synaptic weight | 0 | 1 | |
| $w_{max}$ | Maximum synaptic weight | 1 | 3 | |
| $\tau_{Ca,NMDA}$ (ms) | Nonlin. $Ca^{2+}$ timescale | 80 | 250 | We chose a relatively high lower bound to eliminate interference between the linear and nonlinear calcium transients in the fitting. |
| $\eta$ ($ms^{-1}$) | Nonlinearity parameter | 0 | 500 | |

**Table 4.** Model fit and prediction errors are compared to (a) an estimate of the data variance (3rd column) and to (b) the error of a null model (4th column). Our estimate of the data’s variability is obtained by computing the squared error of every data point relative to the mean of all measurements of the same protocol, averaging over all points within a category of the dataset (pair / burst / high-frequency / imaging), and taking the square-root. For the spike-pair protocols we bin points in terms of Δt and the extracellular calcium so there are at least 2 data points in each bin. The null-model error is computed by assuming no synaptic change under any experimental condition (*Δw* = 1). For the imaging experiments, the null model assumes transient calcium amplitude is independent of the extracellular concentration, and the calcium entry is linear in the number of post-synaptic spikes.

| Error | Meaning | Data variance estimate | Null Model |
| --- | --- | --- | --- |
| $\epsilon_{pair}$ | Pre-post pair error | 0.184 | 0.258 |
| $\epsilon_{burst}$ | Post-synaptic burst error | 0.299 | 0.377 |
| $\epsilon_{total}$ | Total low freq. plasticity error | 0.248 | 0.323 |
| $\epsilon_{freq}$ | High freq. plasticity error | 0.255 | 0.350 |
| $\epsilon_{imaging}$ | Imaging error | 0.534 | 1.282 |

The fitting error is defined to be RMS error over the 144 data points (black crosses in Figure 8A). The fitting error is highly nonlinear in terms of the model parameters, and therefore we cannot expect it to be convex. Hence, for each model we initialized a gradient descent routine at 2000 points chosen uniformly at random within the allowed parameter hypercube. We used the nonlinear constrained optimization built-in to the Matlab software, choosing the active-set algorithm. Using a different optimization algorithm (interior-point) had impact on the time it took to find (local) minima and the fraction of initial conditions of the parameter set that “exited” the allowed hypercube, due to the differences in how step sizes are computed in different algorithms. However, using a different algorithm did not affect the properties of parameter sets with small fitting errors.

#### Parameter variation

To study the effect of varying model parameters in the neighborhood of the best fitting parameter set we multiplied each model parameter by a factor 1 + 0.1x, where x is drawn at random (independently for each parameter and each repetition of this procedure) from a standard normal distribution. With these randomized parameter sets in hand, we computed the STDP curves and the changes in synaptic efficacies following all experimental protocols. These were then used to compute the error bars and shaded areas around the curves showing *Δw* as a function of *Δt* and *Δw* as a function of the pairing-frequency *f*.

#### Eliminating the linear postsynaptic contribution

Previously published results have indicated that repeated post-synaptic stimulation, even at high frequencies, does not lead to long-term synaptic plasticity if the pre-synaptic neuron does not fire ^41, 42^. Qualitatively, the model described above is inconsistent with this observation, since the post-synaptically induced calcium transient alone will summate and cross the LTD/LTP threshold for high enough frequency. We therefore asked whether our model can be modified such that it will predict no change in the synaptic efficacy for protocols where only the post-synaptic neuron is stimulated. Indeed, we find that, since the main contribution to calcium transients is nonlinear (Figure 7), simply dropping the post-synaptic only contribution to the total calcium transient leads to negligible changes to the model predictions for the plasticity experiments (compare Figure 8 with Supplementary Figure 7). Note that the post-synaptic component of the transient is still used for the purposes of computing the nonlinear contribution. We focus our discussion on the “full” model (that includes linear dependence on post-synaptic activity), but we emphasize that the data can be fit equally well with a model with such linear dependence.

---

**Supplementary Figure 1.**
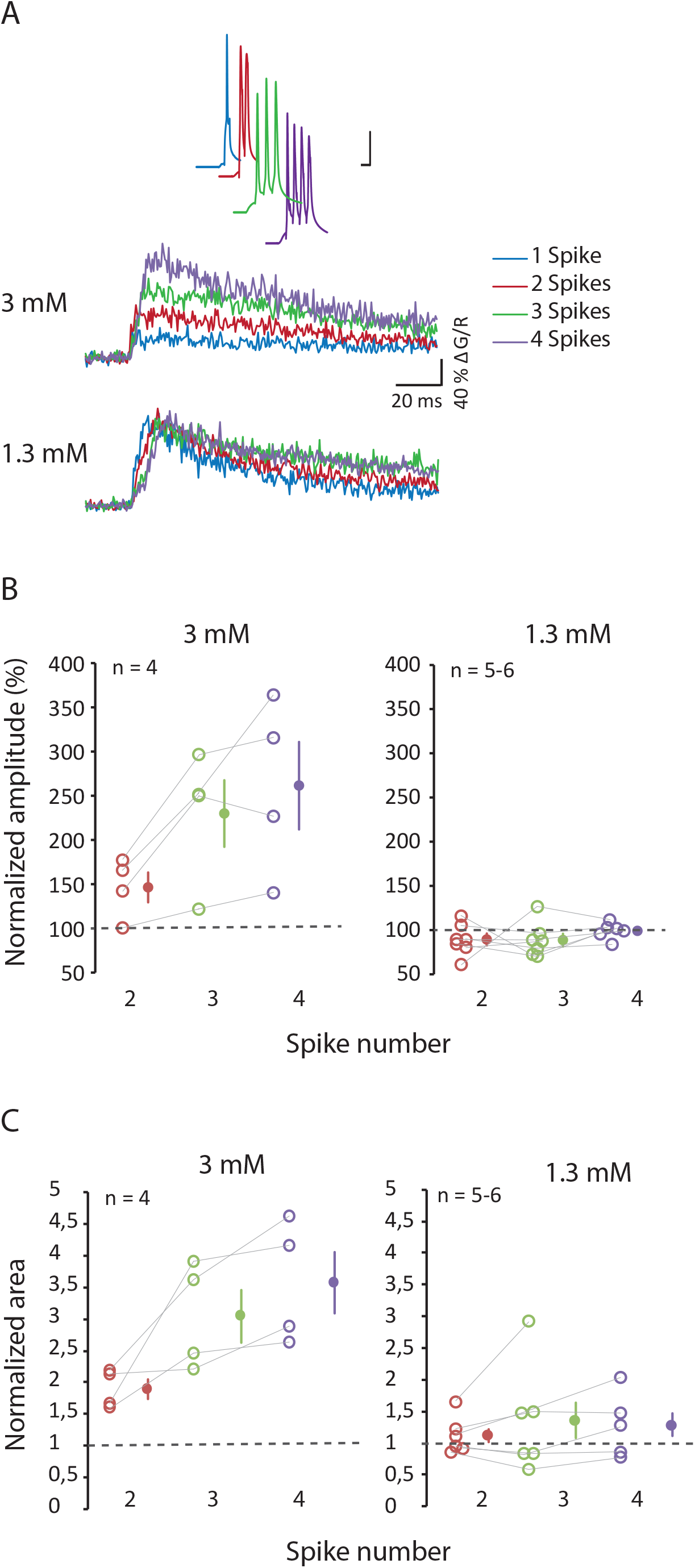
Comparison of calcium transient during at 2 calcium concentrations. (**A**) Representative traces of calcium signals measured in a single dendritic spine evoked by a pre-post protocol at Δt = +20 ms with one (blue traces), two (red traces), three (green traces) or four (purple traces) action potentials. (**B** & **C**). An almost linear summation for amplitude (**B**) and integral (**C**) is observed in 3 mM extracellular calcium but not in 1.3 mM extracellular calcium.

**Supplementary Figure 2.**
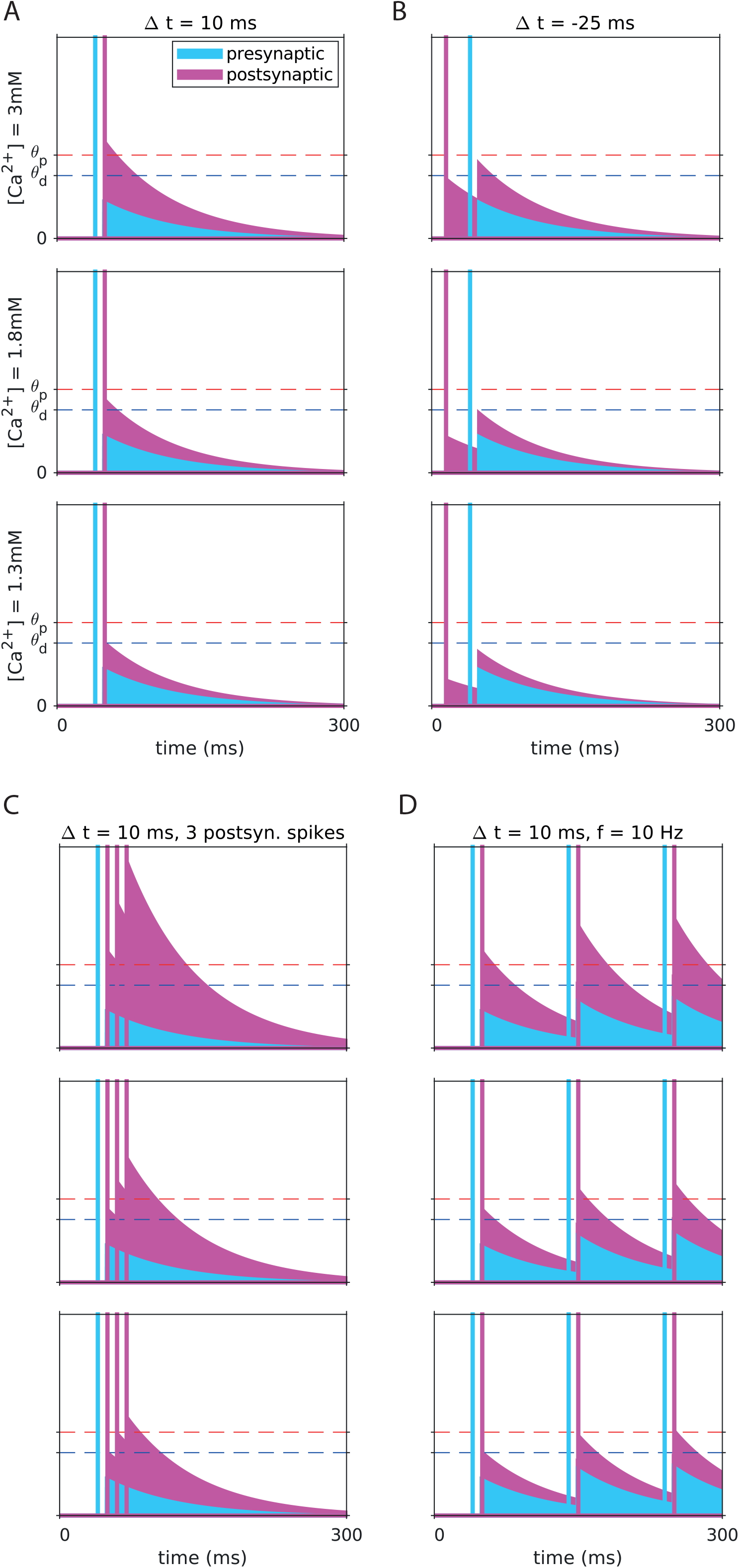
Dependence of calcium transients of linear plasticity model on the stimulation protocol and on the extracellular calcium concentration. Same as Figure 7, for a model with calcium transients that are linearly dependent on pre- and post-synaptic activity. Transients resulting from different protocols qualitatively match the observed plasticity for the same protocol in the experiment.

**Supplementary Figure 3.**
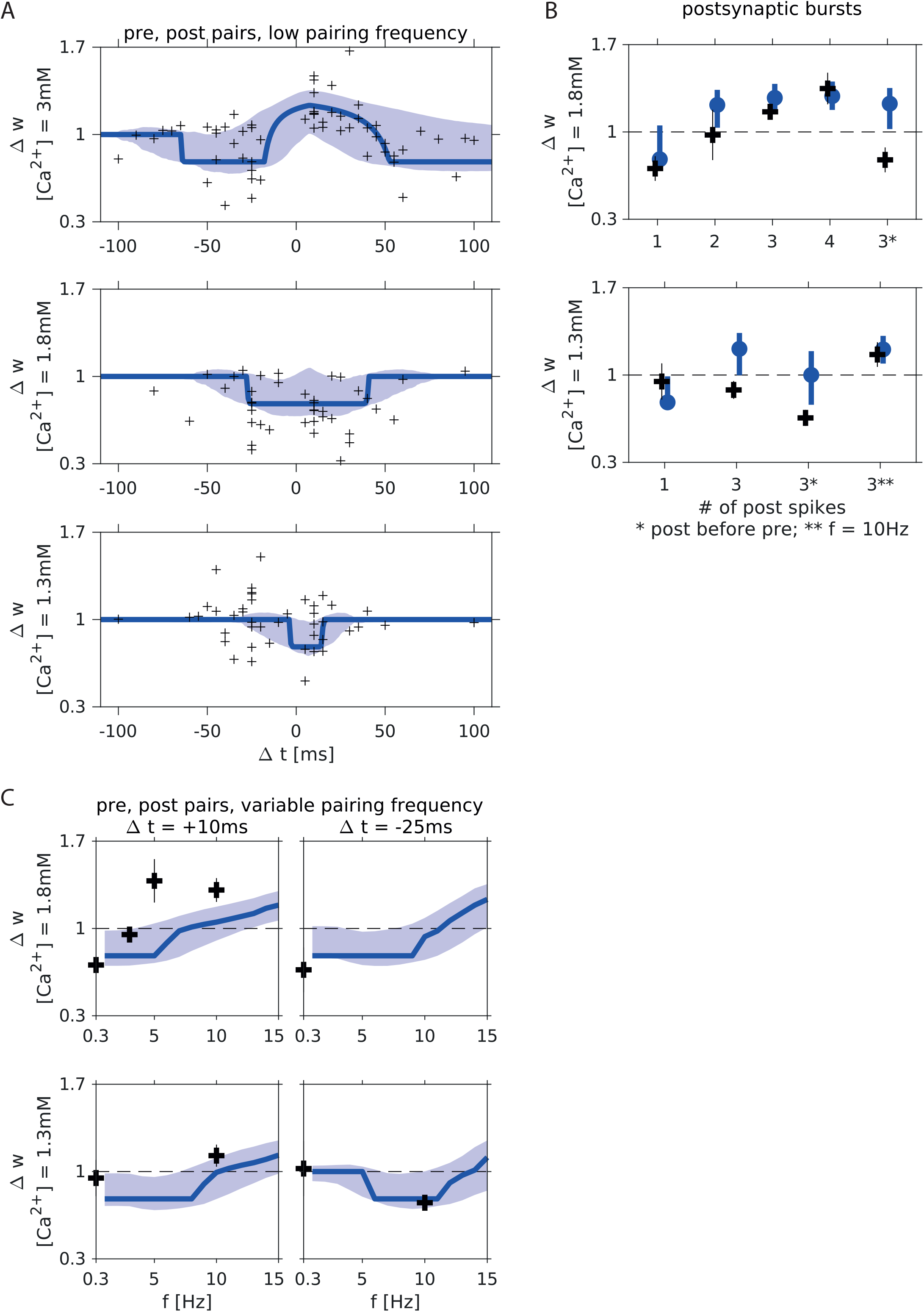
A linear calcium based plasticity model fails to quantitatively account for the plasticity data at multiple concentrations. Same as Figure 8, for a model with calcium transients that are linearly dependent on pre- and post-synaptic activity. For spike pairs at small Δt the model correctly captures the change of sign of plasticity as the extracellular calcium concentration changes. However, the inferred decay timescale of calcium transients is long, so this model predicts significant changes to the synaptic weight even when pre and post-synaptic spikes are separated by more than 100 ms. The long decay timescale also implies saturation to LTP as a function of the number of post-synaptic spikes in burst protocols and as a function of the pairing frequency, leading to poor predictions.

**Supplementary Figure 4.**
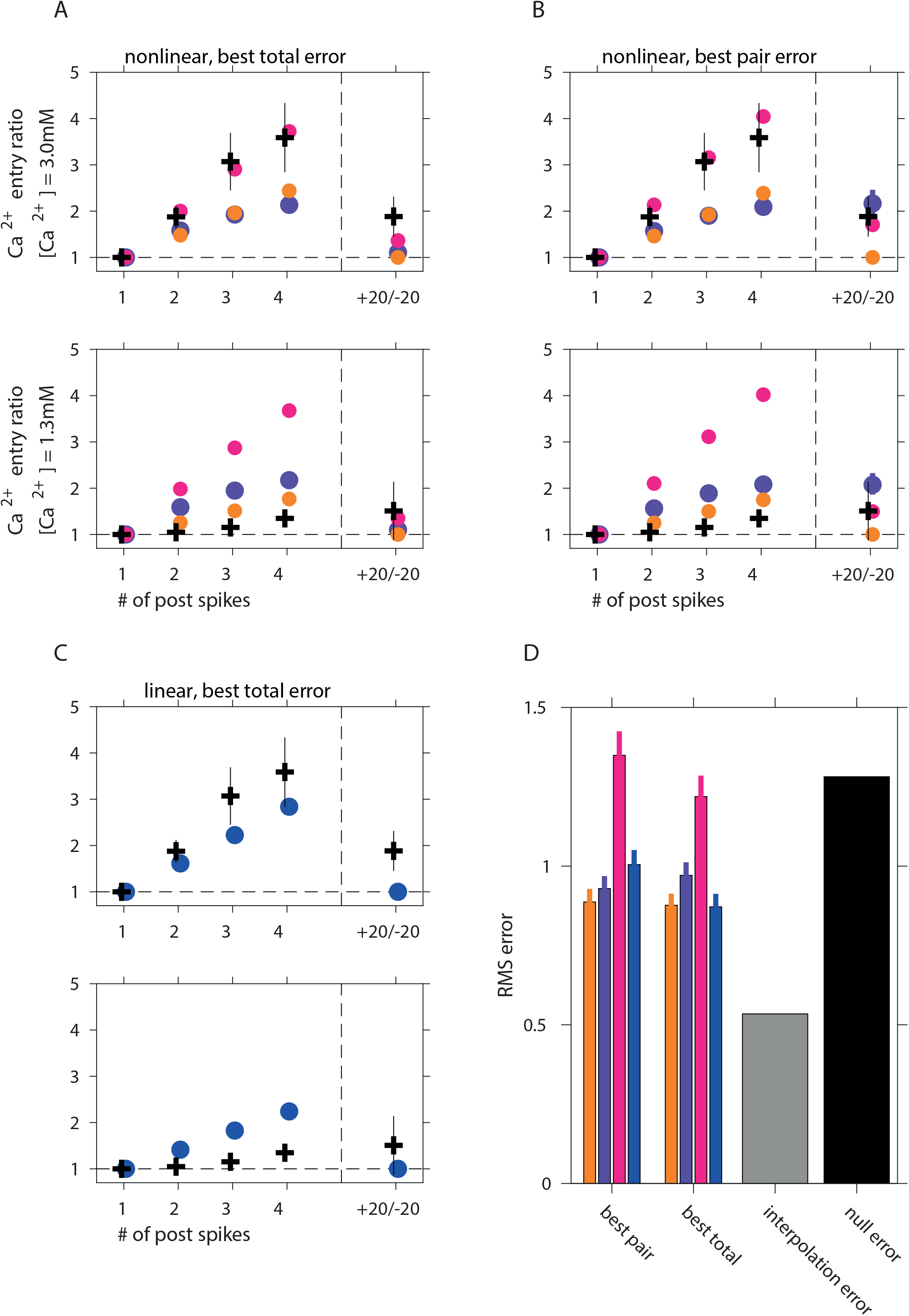
Using calcium imaging of intracellular transients to constrain models. For every model variant shown in the paper we plot the total calcium entry ratio as a function of the number of postsynaptic spikes, the relative timing, and the extracellular concentration. Data from protocols with one and two post-synaptic spikes was used to constrain model parameters (see Methods). Protocols measuring calcium entry as a function of timing (Δt = +20/-20 ms, pre-post vs. post-pre, in the same neurons; see Figure 4 and Supplementary Figure 1) imposed weaker constraints on model parameters, so are shown here despite not being used in the fitting procedure (see Methods). (**A**) Nonlinear models chosen based on lowest combined error for spike-pair and burst plasticity protocols. (**B**) Nonlinear models chosen based on lowest error for spike-pair plasticity protocols. In panels A and B we show results for models fitted subject to constraints imposing varying levels of accuracy relative to the imaging experiments: yellow, 1 standard deviation; purple, 2 standard deviations; red, no constraints from imaging experiments. (**C**) A linear model chosen based on lowest combined error for spike-pair and burst plasticity protocols. (D) Root-mean-square error of calcium entry ratios of each model variant averaged over all data points. Similarly to Figure 8, gray and black bars indicate estimate of the data variability and the errors of a null model, respectively.

**Supplementary Figure 5.**
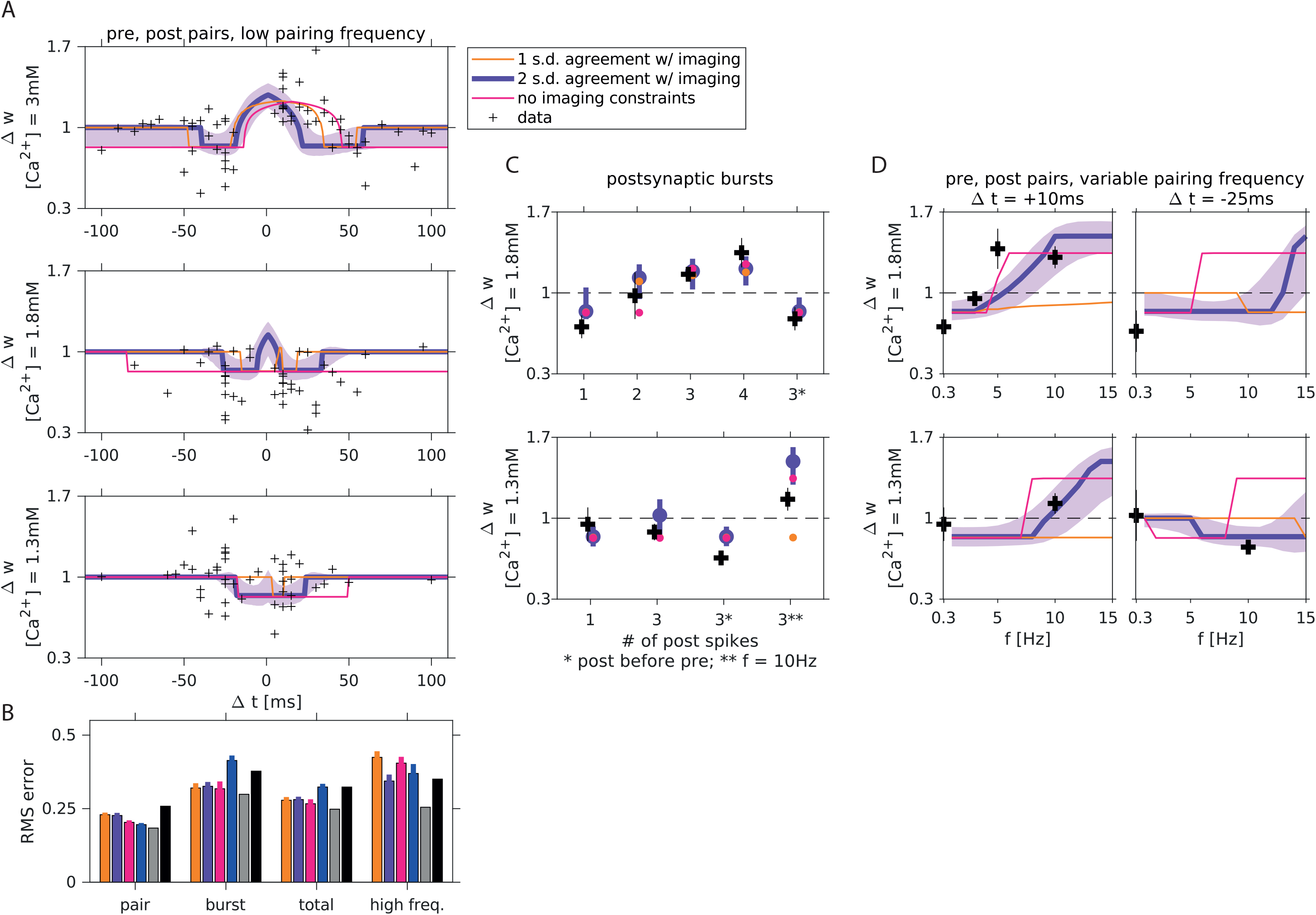
Constraints on parameters of nonlinear model from imaging experiments improve predictions for high pairing frequency experiments. Same as Figure 8, here we also show results for a nonlinear model where parameters were unconstrained by imaging experiments (red) and using strong/weak constraints based on the imaging experiments (orange/purple). Imposing a weak constraint on parameters using the imaging experiments (2 STD of r_2_, see Methods) leads only to a small increase in the fitting error, and to substantial improvements to the predictions for the high-frequency data not used for fitting or model selection. The imaging experiments thus serve as an important regularization on the fitting procedure. Strong constraints by the imaging experiments (1 S.D. of r_2_, see Methods) lead on the other hand to poor fits and predictions to the plasticity experiments, especially protocols with high pairing-frequencies.

**Supplementary Figure 6.**
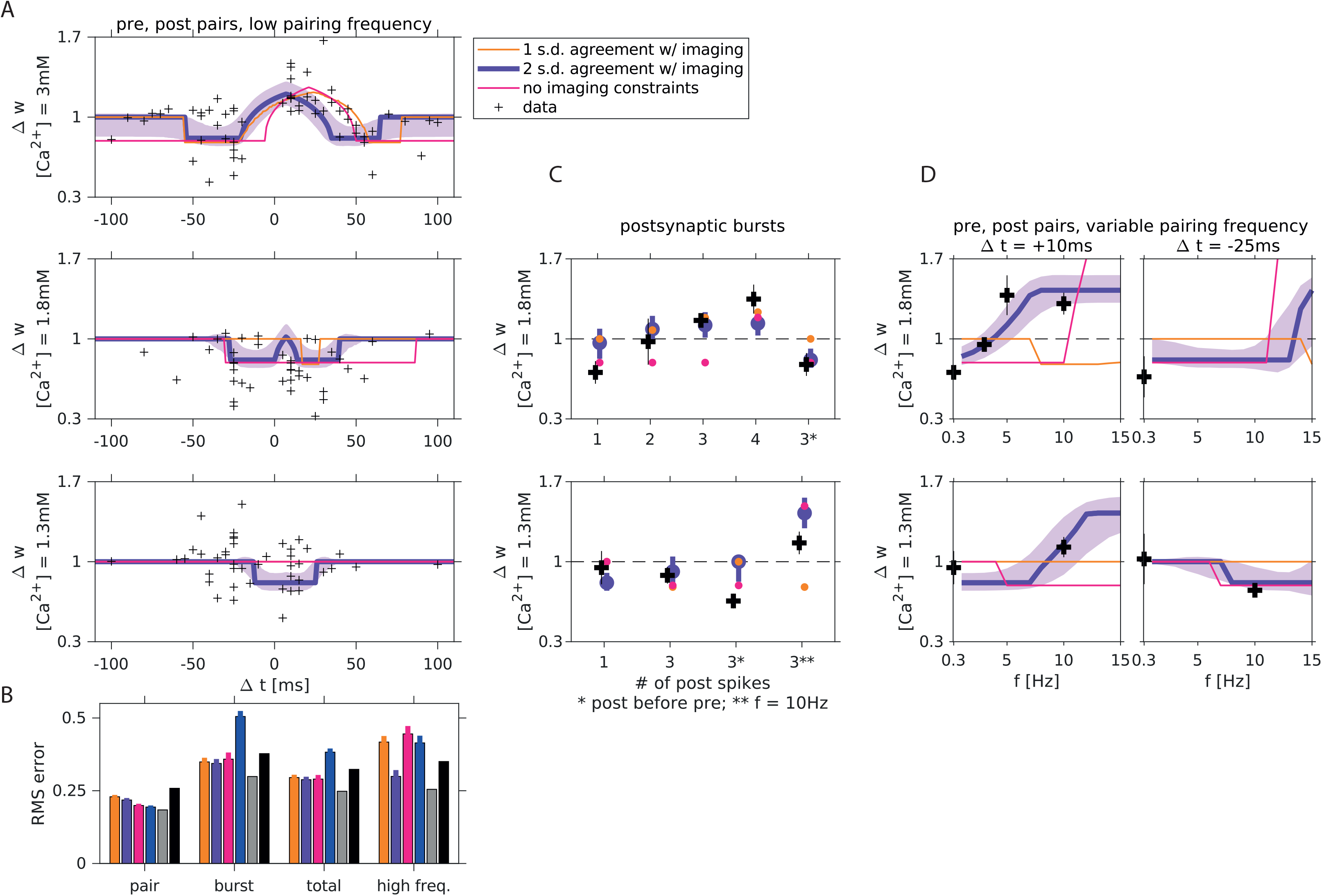
Nonlinear models fit to spike-pair data alone produce accurate predictions. Same as Figure 8, Supplementary Figure 5, here we show results for parameter sets yielding the lowest fit error to spike-pair data (compared to Figure 8 where showing results for parameter sets fit to spike-pair data, but selected to have lowest combined spike-pair and burst data error). As expected the prediction error for burst stimuli is increased, but the model is still qualitatively consistent with the entire dataset. In fact, the predictions to the held-out data (high pairing frequency protocols) is better here compared to Figure 8, suggesting that our model does not suffer from over-fitting.

**Supplementary Figure 7.**
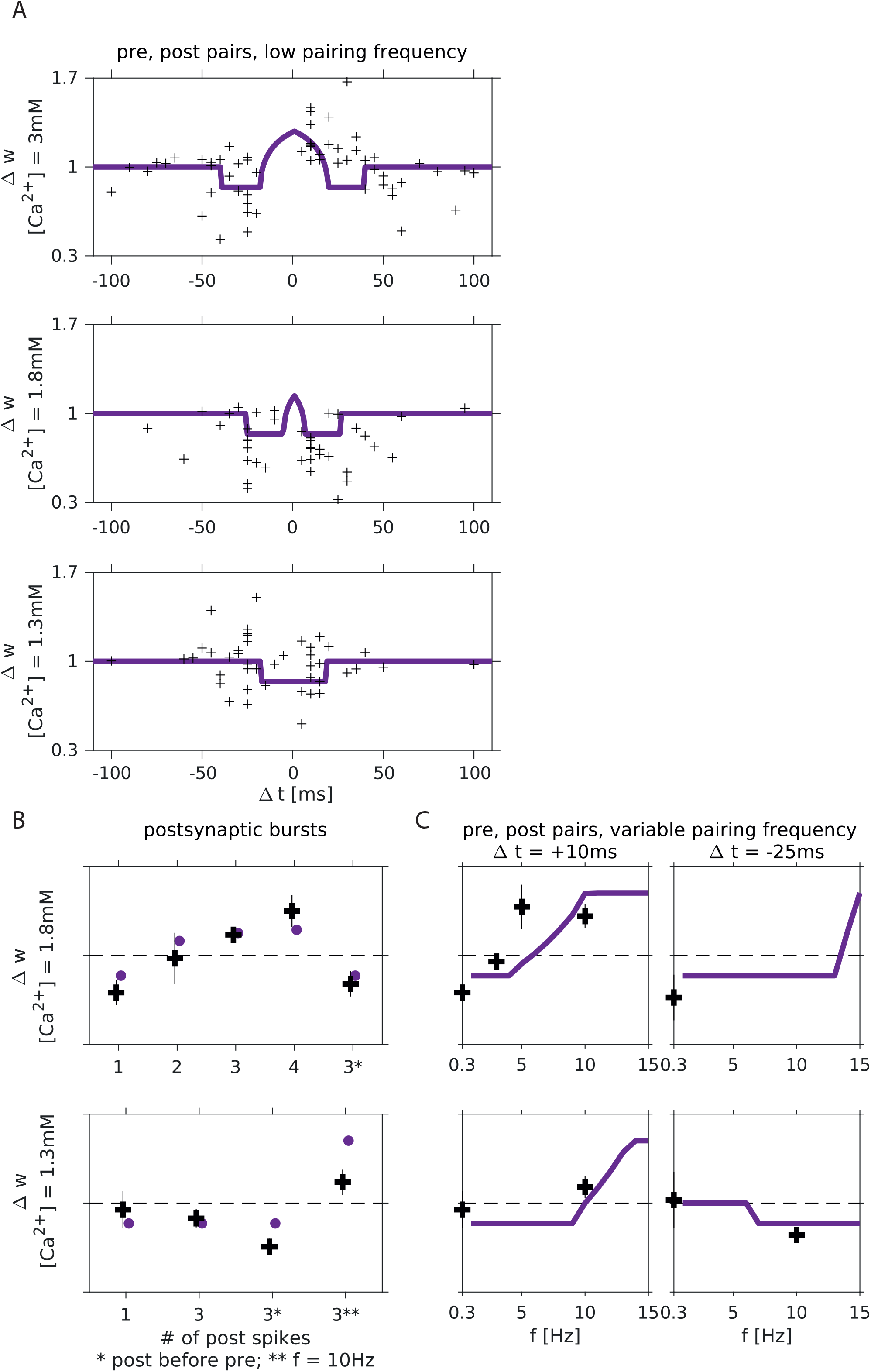
A nonlinear model without direct post-synaptic contribution to calcium transients produces accurate predictions. Same as Figure 8, here we show results for the same model discussed in the main text, where the post-synaptic contribution is eliminated (see Methods). The differences in predictions are negligible. Thus, our model is consistent with experiments showing that repeated post-synaptic stimulation alone, even at high frequencies, does not lead to plasticity.

## Supplementary mathematical note

### 1 Model fitting

We fit the calcium based plasticity model to the experimental STDP curves at three calcium concentrations using the procedure described below. The central part of the fitting procedure is to mathematically express the relative change in synaptic efficacy Δ*w* as a function of the induction protocol, the extra-cellular calcium concentration [Ca^2+^] and the model parameters.

The function Δ*w* is computed in two steps as a function of these variables. First we compute the time the intra-cellular calcium variable *c*(*t*) spends above the depression and potentiation thresholds: *T_d_* and *T_p_,* respectively. In the second step we use *T_d_*, *T_p_* and the rest of the variables and parameters to compute Δ*w*.

### Computation of calcium transients and time spent above threshold

The calcium transient *c*(*t*) is a sum of three contributions: pre-synaptic, post-synaptic and nonlinear, unless noted otherwise. The contributions to the total calcium transient evolve according to the differential equations:

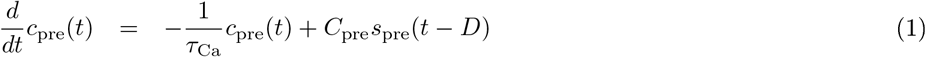

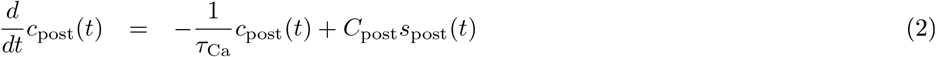

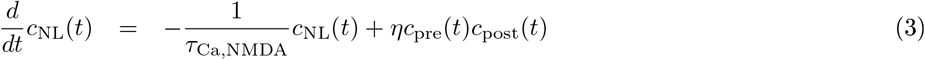

Where *C*_pre_, *C*_post_ are the pre- and post-synaptic amplitude parameters, 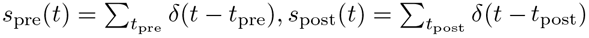 the pre- and post-synaptic spike trains (i.e., sums over Dirac-delta functions centered at the spike times), *D* is the delay parameter and *η* is the nonlinearity parameter.

For induction protocols with low pairing frequencies one can assume that the transient calcium returns to baseline before each repetition. This assumption holds for a paring frequency of 0.3Hz used for most of our measurements (and *all* the data used for model fitting purposes), making it sufficient to compute transients for a single repetition of each protocol. Furthermore, when a protocol consists of a single pre- and a single post-synaptic spike occurring at *t* = 0 *t* = *Δt*, respectively, Eqs. (1-3) can be solved analytically:

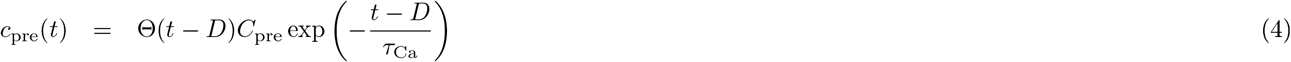

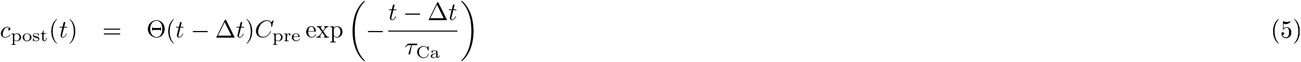

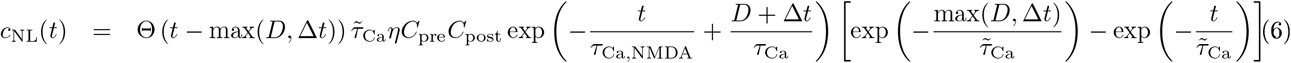

Here ϴ(-) is the Heaviside step function and 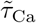 is the effective time constant:

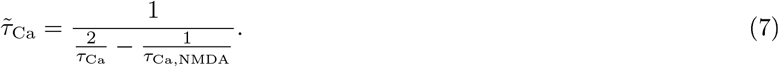

The calcium transient was computed using these expressions:

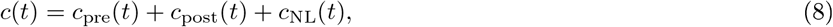

with temporal resolution *dt* = 0.25 ms.

For protocols with a post-synaptic burst of spikes, *c*_post_ and *c*_NL_ were computed by replacing Δ*t* in Eqs (5,6) with the times of the spikes in the post-synaptic burst (relative to the pre-synaptic spike at *t* = 0) and summing over all the spikes in the burst of the post-synaptic neuron^1^.

For protocols with high pairing frequency where the calcium transient does not necessarily return to baseline following each repeat of the induction protocol, the transients were computed using the Euler method with time-step *dt* = 0.25ms. Using a smaller time-steps made fitting considerably slower but it did not change the results.

The time spent above the depression and potentiation thresholds (during each repeat of the induction protocol of duration *T*) is then

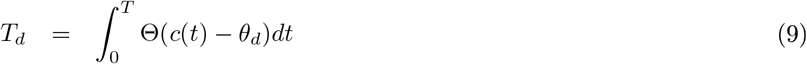

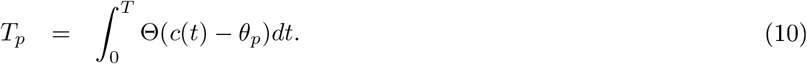

The integrals were approximated by counting temporal bins in which *c*(*t*) exceeded the threshold and multiplying by *dt*. Note that for the linear model (*η* = 0, *T_d_* and *T_p_* can be computed analytically (see Graupner and Brunel, 2012).

### Computation of *Δw*

Recall that in our model, the synaptic weight variable evolves according to

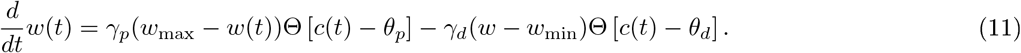

From this, one can show that if the initial value *w*(*t* = 0) = 1 and the protocol is repeated *n* times, the synaptic weight variable at the end of the protocol *Wf* = *w*(*t* = *nT*) is

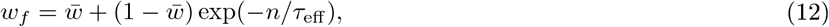

where

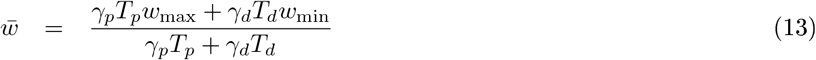

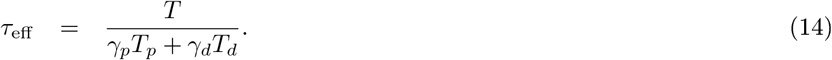

By assumption the synaptic weight at time 0 is *w*(*t*) = 1, so the relative change in synaptic efficacy as a function of the protocol and the model parameters is simply

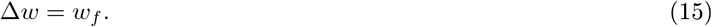

### 2 Constraints from imaging experiments on model parameters

#### 2.1 Linear model

For the linear model, we analytically derive constraints on the model parameters *a*_post_, *a*_pre_, *C*_post_, *C*_pre_ given the results of the imaging experiments. Let *I*_1_ be the total calcium entry resulting from a single pre- and a single postsynaptic spike. We have,

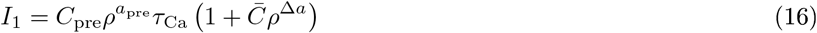

where *ρ* is the extracellular calcium concent ration, and 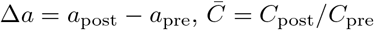

Similarly, let *I*_2_ be the total calcium entry resulting from a single pre- and two postsynaptic spikes. Now we have,

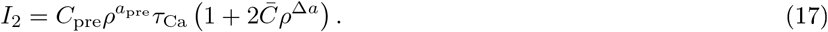

Define r_2_(*ρ*) to betheratiooftotalcalciumentry,

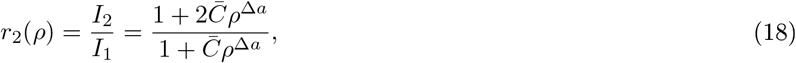

So,

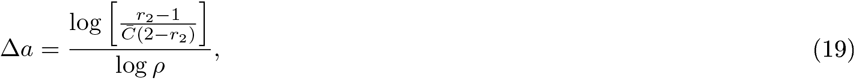

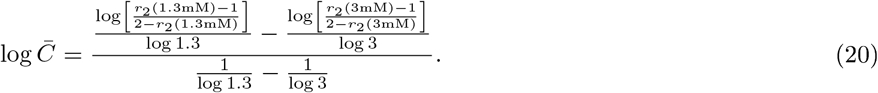

From the experimental data we have

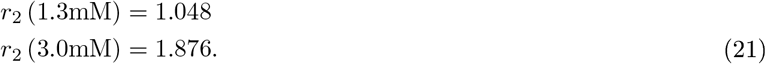

giving

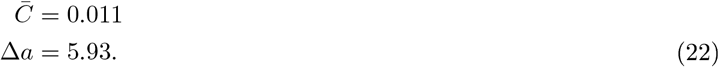

#### 2.2 Nonlinear model

To express *r*_2_ as a function of the full model parameters (including the nonlinearity), we compute the integral over the pre, postsynaptic and nonlinear terms [Eqs. (4-6)]. Restoring the explicit dependence on the calcium concentration *ρ*,

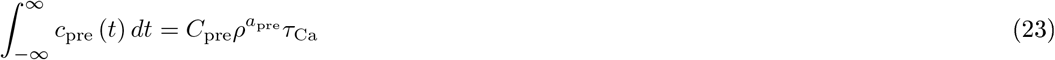

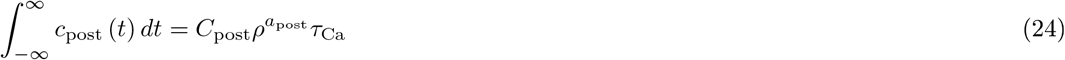

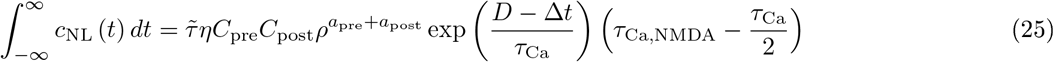

Using these and the experimentally measured values of *r*_2_ we added inequalities as constraints to our numerical optimization procedure. Specifically, the model values of *r*_2_ were required to be within either 1 or 2 standard devaitions of the empirical measurement.

## Footnotes

1 This is only valid because there was a single pre-synaptic stimulation in every repetition of the induction protocol. Had there been more than a single pre-synaptic stimulation then computing *c*_NL_ requires solving Eqs. (4-6) explicitly.

